# Synapses and Ca^2+^ activity in oligodendrocyte precursor cells can predict where myelin sheaths form

**DOI:** 10.1101/2022.03.18.484955

**Authors:** Jiaxing Li, Tania Miramontes, Tim Czopka, Kelly R. Monk

## Abstract

In the nervous system, only one type of neuron-glial synapse is known to exist: that between neurons and oligodendrocyte precursor cells (OPCs). Neuron-OPC synapses are thought to bridge neuronal activity to OPCs. However, their composition, assembly, downstream signaling, and *in vivo* functions remain largely unclear. Here, we use zebrafish to address these questions and identify postsynaptic molecules PSD-95 and Gephyrin in OPCs. They increase during early development and decrease upon OPC differentiation. PSD-95 and Gephyrin in OPCs are highly dynamic and frequently assemble at “hotspots.” Gephyrin hotspots and synapse-associated Ca^2+^ activity in OPCs predict where a subset of myelin sheaths form in oligodendrocytes. Further analyses reveal that spontaneous synaptic release is integral to OPC Ca^2+^ activity, while evoked synaptic release contributes only in early development. Finally, disruption of the synaptic genes *dlg4a&b*, *gephyrinb*, and *nlgn3b* impairs OPC differentiation and myelination. Together, we propose that neuron-OPC synapses are dynamically assembled and can predetermine myelination patterns through Ca^2+^ signaling.

## INTRODUCTION

Oligodendrocyte precursor cells (OPCs) are a dynamic cell type in the central nervous system (CNS) that can migrate, proliferate, dramatically remodel processes, and differentiate into myelinating oligodendrocytes. OPCs and their interactions with neurons play important roles in CNS health and disease such as myelination and remyelination^1^, neural network remodeling^2^, and glioma progression^3^. Synapses form between neurons (presynaptic) and OPCs (postsynaptic) and represent the only type of glial synapse in the nervous system. The composition of neuron-OPC synapses is largely unknown, with the exception of some synaptic receptors (*e.g*., glutamatergic and GABAergic) that were revealed by pharmacology and electrophysiology^4^. Neuron-OPC synapse assembly and regulation *in vivo* are mysterious, due in part to a paucity of tools to label these structures. Previous studies knocking down single synaptic receptor types found diverse and sometimes conflicting effects on OPC development and myelination; for example, blocking NMDA receptors reduced or had no effect on myelination^5,6^, and blocking GABA receptors enhanced or had no effect on OPC proliferation^7,8^. This is possibly due to redundancy, as OPCs possess multiple synaptic receptors^4^. Currently our understanding of the functions of neuron-OPC synapses is incomplete.

OPCs are hypothesized to sense the activity of neurons through neuron-OPC synapses. Several studies support this idea: neuronal activity can modulate OPC development and myelination^9–13^; axonal stimulation in *ex vivo* rodent studies was reported to cause synapse-dependent Ca^2+^ increases in oligodendrocyte lineage cells, mostly in somas^14–16^. However, the *in vivo* relationship between neuronal activity, neuron-OPC synapses, and OPC Ca^2+^ activity without exogenous stimulation is unknown. Furthermore, the role of synapses in forming microdomain (MD) Ca^2+^ events in OPC processes^17,18^ has not been addressed.

Here we identify postsynaptic scaffold proteins membrane-associated guanylate kinases (MAGUKs) and Gephyrin in zebrafish OPCs. The postsynaptic puncta number in OPCs increases during early development, reduces upon OPC differentiation, and exhibits small differences across CNS regions but no differences between OPC subgroups. *In vivo* imaging of PSD-95-GFP (a MAGUK member) and GFP-Gephyrin puncta in OPCs reveals that the puncta are highly dynamic, presumably resulting in extensive synaptic coverage. Although dynamic, postsynaptic puncta repeatedly assemble at approximately 10 hotspots per OPC. GFP-Gephyrin hotspots at the OPC stage predict where a subset of myelin sheaths form at the oligodendrocyte stage, in a synaptic release-dependent manner. Further analysis reveals that Ca^2+^ activity is associated with neuron-OPC synapses and similarly predicts where myelin sheaths form. The majority of *in vivo* OPC Ca^2+^ events occur in MDs, require spontaneous synaptic release, and, during early development, require evoked synaptic release from neurons. Finally, we find that disruption of synaptic genes, *dlg4a*&*b*, *gephyrinb*, and *neuroligin3b* (*nlgn3b*) impairs OPC development and myelination. Collectively, these results suggest that neuron-OPC synapses partially predetermine myelination patterns through preferential assembly and Ca^2+^ signaling.

## RESULTS

### Postsynaptic organizers MAGUK/PSD-95 and Gephyrin are present in zebrafish spinal cord OPCs

Transcript expression of *Dlg4* (encodes PSD-95, often associated with excitatory Glutamate receptors) and *Gephyrin* (often associated with inhibitory GABA and Glycine receptors) were previously reported in zebrafish and mammalian OPCs (Extended Data Fig. 1a-b)^17,19^. We detected the presence of MAGUK/PSD-95 and Gephyrin puncta in zebrafish spinal cord OPCs using immunohistochemistry (IHC) at 3 days post-fertilization (dpf) (Fig. 1a-c and Extended Data Fig. 1c) and found that puncta were localized to OPC processes and somas (Extended Data Fig. 1d). Next, we visualized postsynaptic puncta using Fibronectin intrabodies generated by mRNA display (FingRs) that bind to endogenous PSD-95 and Gephyrin*in vivo*^20^. We found FingR.GFP enrichment in somas (Fig. 1d), similar to previous reports in neurons^20–22^. More importantly, we observed puncta labeling in OPC processes (Fig. 1d), indicating potential synaptic locations. By IHC and FingRs, we identified 18.2 ± 2.5 and 20.3 ± 2.4 MAGUK/PSD-95 puncta, respectively, and 29.2 ± 3.1 and 21.1 ± 4.0 Gephyrin puncta, respectively, per OPC at 3 dpf (Fig. 1e). The similar results from two distinct methods support the presence of postsynaptic puncta in OPCs and extend previous work that reported PSD-95 in myelin sheaths^23^.

**Fig. 1.**
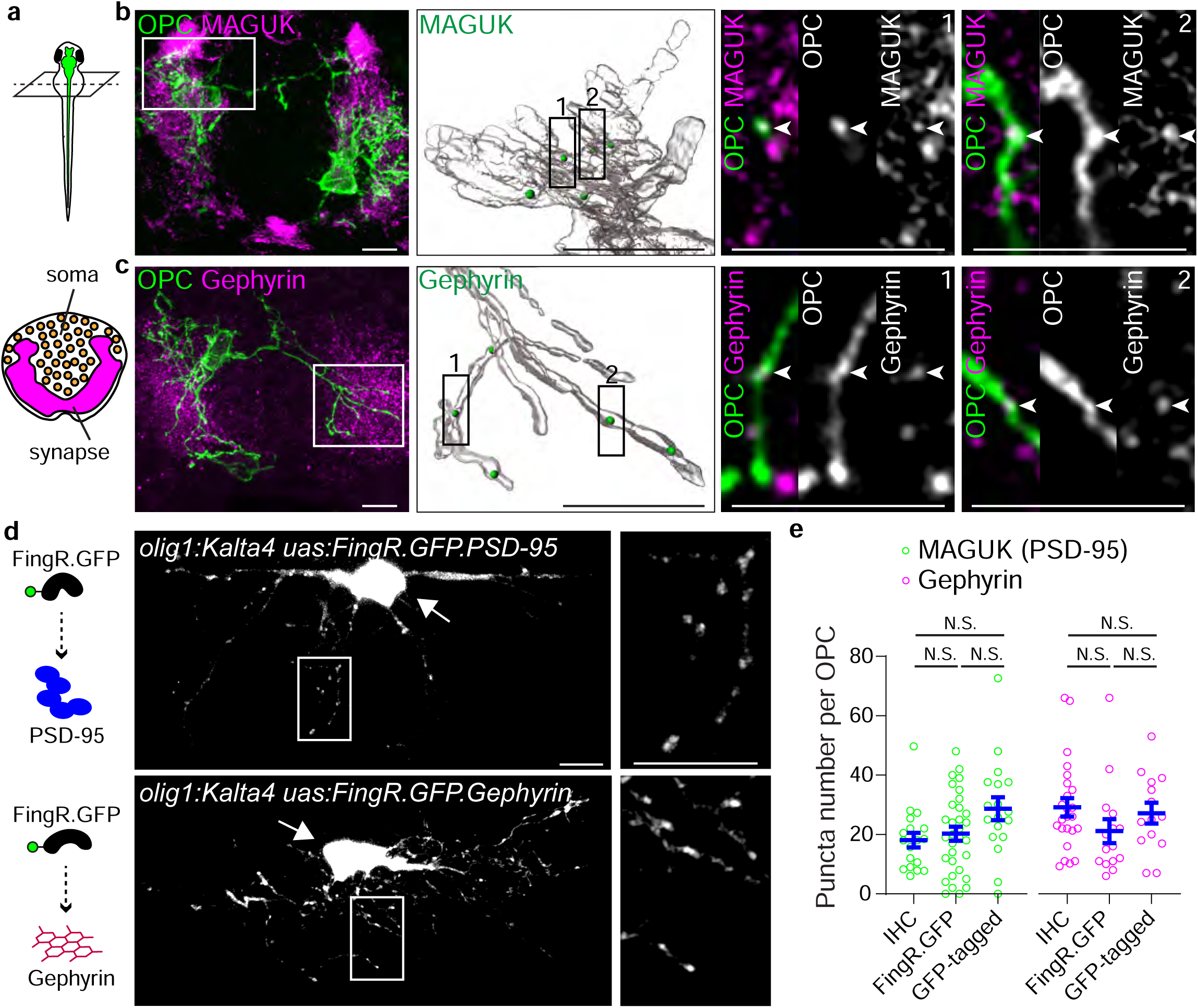
Postsynaptic organizers MAGUK/PSD-95 and Gephyrin are present in zebrafish spinal cord OPCs. (a) Top, schematic fish model showing transverse section orientation; bottom, distribution of soma and synapses in spinal cord transverse sections. (b-c) Representative IHC images of (b) MAGUK and (c) Gephyrin in *Tg(olig1:Kalta4,10xUAS:myrGFP)* spinal cord at 3 dpf. From left, spinal cord sections, 3D-reconstructed regions, and two examples of puncta in OPCs at single planes. Only the postsynaptic puncta that fall within OPC process volumes are shown in 3D-reconstructed images (green spheres) and are indicated with arrowheads in examples. In each example, from left are the merged channels, the OPC channel, and the MAGUK/Gephyrin channel. (d) Representative images and enlarged regions of FingR.GFP.PSD-95 and FingR.GFP.Gephyrin driven by *olig1* promoter in OPCs. OPC somas are indicated with arrows. The process regions are enlarged in the right panel. The schematic on the left shows FingR.GFP binding to the proteins of interest. (e) Quantification of puncta number per OPC by IHC (MAGUK and Gephyrin), FingR.GFP (PSD-95 and Gephyrin), and GFP-tagged (PSD-95 and Gephyrin) in the spinal cord. From left, n = 18, 32, 18, 24, 15, and 14 cells from at least 10 fish each condition. All data are represented as mean ± SEM; (e) Kruskal-Wallis test followed by Dunn’s multiple comparison test; scale bars 5 μm.

Given that OPCs are heterogeneous according to transcriptomic and electrophysiological studies^24,25^, we examined the number of postsynaptic puncta across developmental stages and CNS regions using IHC. We observed an increasing trend through development: both MAGUK and Gephyrin puncta numbers increased from 2 dpf (10.4 ± 1.8 MAGUK; 15.9 ± 2.1 Gephyrin) to 5 dpf (28.8 ± 8.6 MAGUK; 68.2 ± 6.7 Gephyrin) (Extended Data Fig. 2a). The postsynaptic puncta numbers per cell were overall similar across CNS regions, except that Gephyrin puncta numbers were higher in the forebrain and midbrain (20.2 ± 1.2) compared to the spinal cord (13.8 ± 0.9) (Extended Data Fig. 2b), possibly suggesting a preference of OPCs to form synapses with inhibitory neurons in the brain. Within the zebrafish spinal cord, OPC subgroups were previously reported to reside in neuron-rich vs. axon and synapse-rich regions (Extended Data Fig. 2c)^17^. We found that postsynaptic puncta numbers were similar between these OPC subgroups (Extended Data Fig. 2d).

### Some but not all MAGUK and Gephyrin puncta in OPCs are associated with other synaptic molecules

Next, we tested whether these puncta align with presynaptic structures. Using the pan-presynaptic marker Synapsin (Fig. 2a-f), we identified MAGUK and Gephyrin puncta in OPCs that aligned (Fig. 2b, e) or that did not align (Fig. 2c, f) with Synapsin. 46.1 ± 7.4% and 38.3 ± 4.8% of MAGUK and Gephyrin puncta in an OPC, respectively, aligned with Synapsin in the spinal cord, and a similar percentage of puncta in an OPC aligned with Synapsin in other CNS regions (Fig. 2g). In addition, at 3 dpf, 42.1 ± 4.8% of Gephyrin puncta in an OPC aligned with Glycine receptors (Fig. 2h-i). We noted that postsynaptic markers in individual OPCs exhibited a wide range of alignment with presynaptic molecules (0-100%). Altogether, the alignment of MAGUK and Gephyrin puncta with other synaptic molecules varies from OPC to OPC, and averages 40-60% throughout the CNS.

**Fig. 2.**
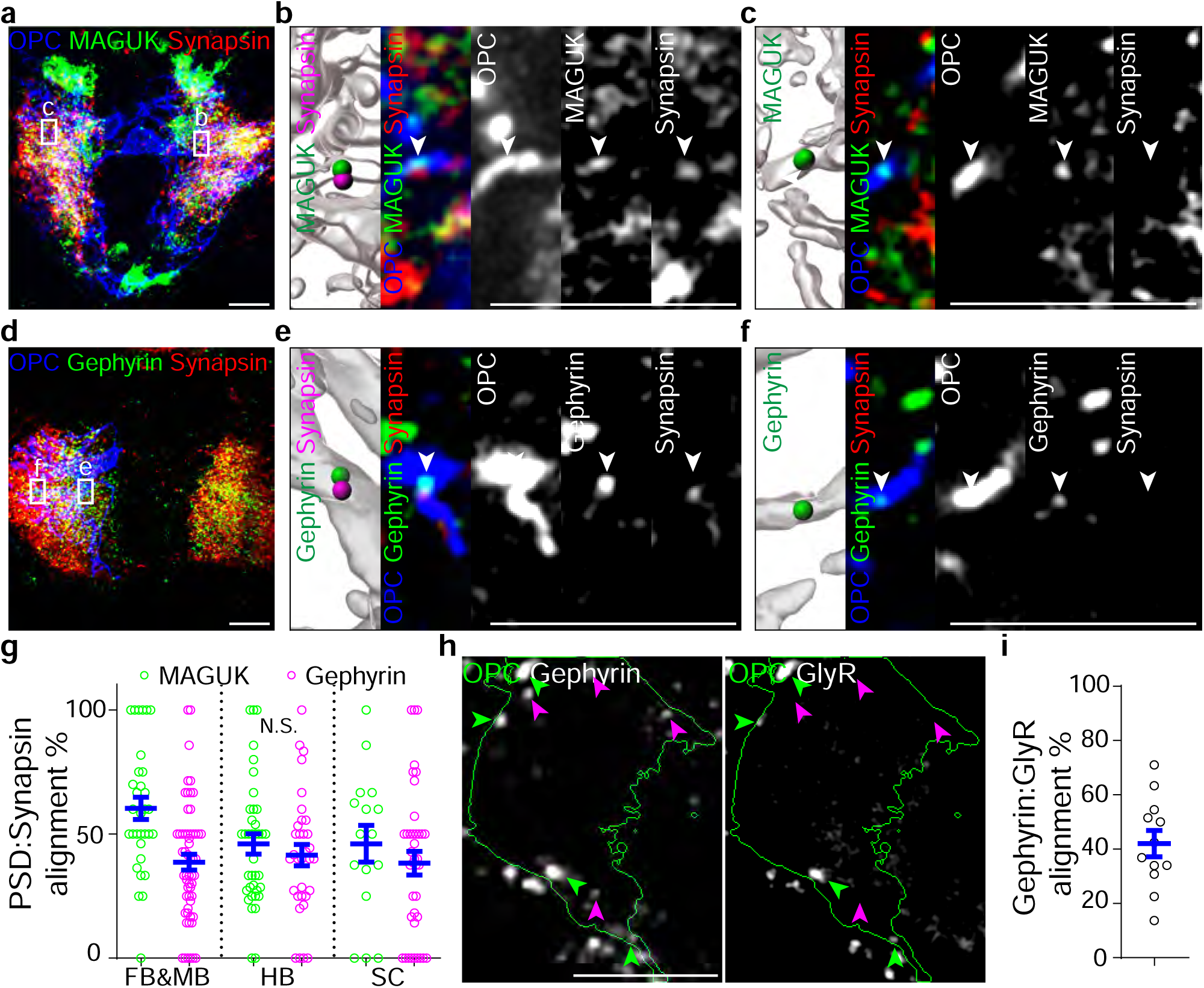
Some but not all MAGUK and Gephyrin puncta in OPCs are associated with other synaptic molecules. (a-f) Representative transverse IHC images of (a-c) MAGUK and (d-f) Gephyrin with Synapsin in *Tg(olig1:Kalta4,10xUAS:myrGFP)* spinal cord OPCs at 3 dpf. (a, d) spinal cord sections, (b, e) alignment example with Synapsin, (c, f) no-alignment example with Synapsin. In each example, from left are 3D reconstructed regions, the merged channels, the OPC channel, the MAGUK/Gephyrin channel, and the Synapsin channel. Only the postsynaptic puncta that fall within OPC process volumes are shown in 3D-reconstructed images (green spheres) with aligned synapsin (magenta spheres) and are indicated with arrowheads in other channels. (g) The % of MAGUK and gephyrin puncta in an OPC that align with presynaptic Synapsin in different CNS regions at 3 dpf. FB&MB, forebrain and midbrain; HB, hindbrain; SC, spinal cord. From left, n= 32, 57, 38, 33, 16 and 39 cells from at least 10 fish each condition. (h) Representative IHC images of Gephyrin and Glycine receptors (GlyR) in *Tg(olig1:Kalta4,10xUAS:myrGFP)* spinal cord OPCs at 3 dpf. OPC boundaries are indicated in green. Gephyrin puncta either align with GlyR (green arrowheads) or do not (magenta arrowheads). (i) The % of Gephyrin puncta in an OPC that align with GlyR in spinal cord at 3 dpf. n = 12 cells from 7 fish. All data are represented as mean ± SEM; N.S., not significant; (g) Kruskal-Wallis test followed by Dunn’s multiple comparison test; scale bars 5 μm.

### *In vivo* imaging reveals downregulation of PSD-95-GFP and GFP-Gephyrin upon OPC differentiation

To understand synapse assembly and disassembly in OPCs, we conducted*in vivo* time-lapse imaging. We found that FingR.GFP tools were especially susceptible to photobleaching during long-term imaging; we therefore generated PSD-95-GFP and GFP-Gephyrin constructs and drove their expression with *sox10*, a pan-oligodendrocyte lineage promoter that also labels a subset of interneurons. At 3 dpf in the spinal cord, we found both markers present in punctate and cytosolic forms in OPCs (Fig. 3a), consistent with patterns observed in neurons^26,27^. PSD-95-GFP and GFP-Gephyrin puncta size in OPCs resembled that of neurons (Fig. 3a-b and Extended Data Fig. 3a). GFP was enriched in somas, a known phenomenon observed with GFP-tagging and knock-in synaptic labeling^28–35^. It is difficult to resolve puncta with such enrichment; therefore, we excluded somas from our analyses. We conducted a series of analyses including puncta number quantification, correlation between puncta number and GFP-tagged protein levels, synaptic alignment, developmental changes, and GFP colocalization with antibody staining (Fig. 1e, 2g and Extended Data Fig. 2a, 3b-i). By every applicable metric, we found that PSD-95-GFP and GFP-Gephyrin are similar to endogenous labeling by IHC and FingR.GFP. Next, we examined if these puncta were regulated upon OPC differentiation. We tracked individual OPCs over the course of their development and quantified the GFP-tagged puncta number. Because OPC differentiation takes several hours for all processes to either retract or transform into nascent myelin sheaths^36,37^, we used the disappearance of the last exploratory process as a reference point. We found a decreasing trend during OPC differentiation: puncta numbers reduced from 9.8 ± 2.5 PSD-95-GFP and 16.5 ± 2.8 GFP-Gephyrin to 3.9 ± 1.2 PSD-95-GFP and 9.1 ± 2.0 GFP-Gephyrin (Fig. 3c-d). In cases where we observed differentiating oligodendrocytes that possessed both processes and myelin sheaths, PSD-95-GFP and GFP-Gephyrin puncta were 4-fold and 3.3-fold more enriched in processes than in sheaths, respectively (Fig. 3e-f), suggesting preferential localization of synaptic puncta to processes rather than nascent myelin sheaths. Altogether, these results show a downregulation of GFP-tagged postsynaptic puncta upon OPC differentiation, consistent with reduced synaptic inputs reported from*ex vivo* electrophysiological studies in rodents^38,39^.

**Fig. 3.**
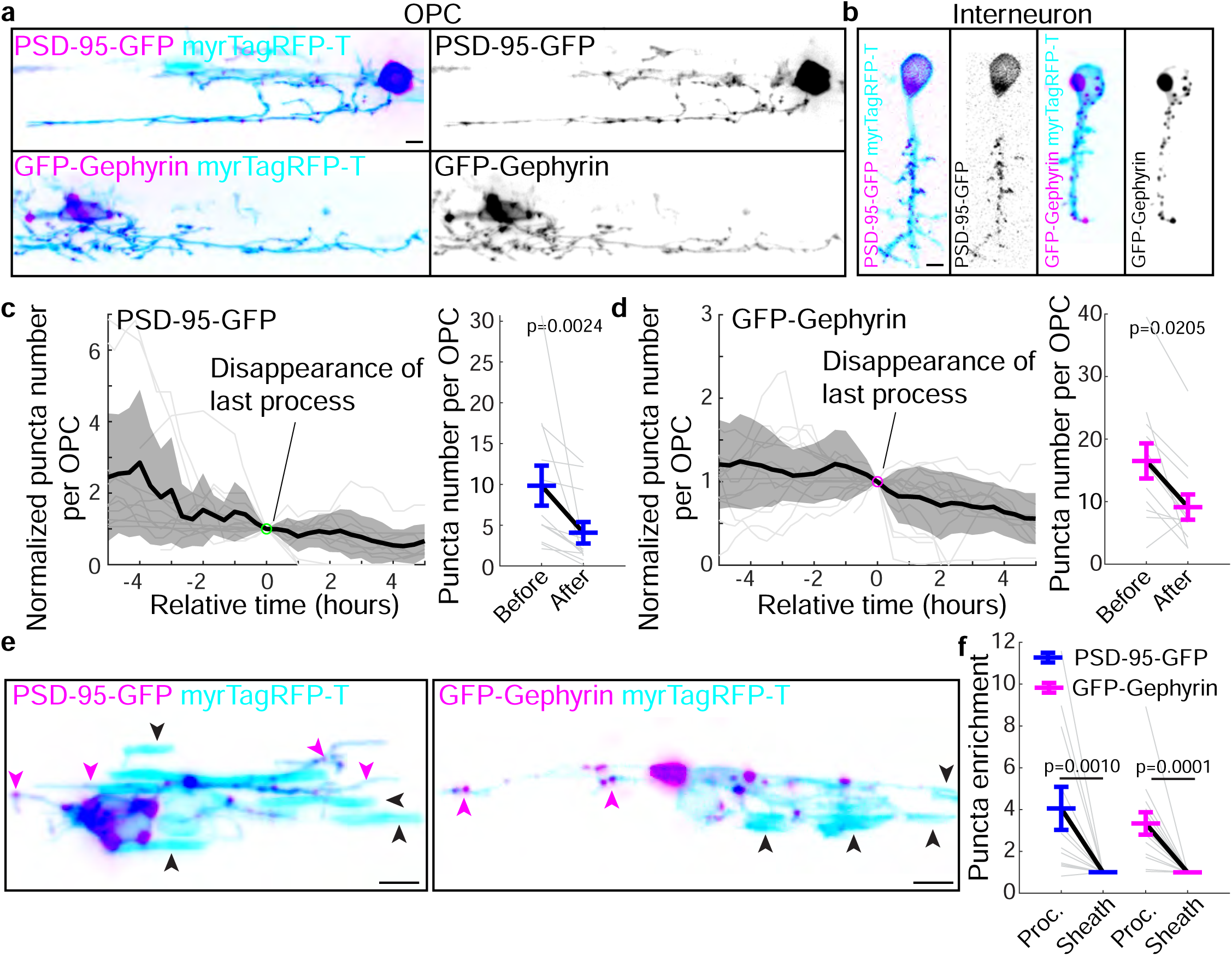
*In vivo* imaging reveals downregulation of PSD-95-GFP and GFP-Gephyrin upon OPC differentiation. (a-b) Representative images of PSD-95-GFP and GFP-Gephyrin (magenta) in spinal cord (a) OPCs and (b) neurons at 3 dpf. Cell membranes indicated with cyan. (c-d) Quantification of (c) PSD-95-GFP and (d) GFP-Gephyrin puncta number in OPCs before and after the last process disappearance, set as time zero, during differentiation. Left, individual traces (grey lines) with the average (black line) and the standard deviation (shade). Right, paired comparison before and after time zero. (c) n= 12 cells from 8 fish; t[11] = 3.922. (d) n= 12 cells from 8 fish; t[11] = 2.705. (e) Representative images of PSD-95-GFP and GFP-Gephyrin (magenta) in differentiating oligodendrocytes (cyan) at 2-3 dpf. Nascent sheaths are indicated with black arrowheads, and processes are indicated with magenta arrowheads. (f) Enrichment of PSD-95-GFP and GFP-Gephyrin puncta in processes and sheaths of differentiating oligodendrocytes. n = 12 and 14 from at least 10 fish each condition. All data are represented as mean ± SEM; N.S., not significant; (c,d) ratio-paired *t*-test; (f) Wilcoxon matched-pairs signed rank test; scale bars 5 μm.

### PSD-95-GFP and GFP-Gephyrin puncta in OPCs are dynamic and repeatedly assemble at hotspots

We next sought to analyze the dynamics of PSD-95-GFP and GFP-Gephyrin in OPCs*in vivo*. These GFP-tagged puncta in OPCs were dynamically assembled and disassembled during an 11-hour imaging session, while those in neurons remained relatively stable (Fig. 4a and Supplementary Video 1). The number of puncta per cell remained similar in neurons, but fluctuated in OPCs over 30 minutes of observation (Fig. 4b). Individual GFP-tagged puncta in neurons lasted about 234 min (PSD-95-GFP) and 206 min (GFP-Gephyrin), while those in OPCs lasted about 12 min (PSD-95-GFP) and 5 min (GFP-Gephyrin) (Fig. 4c). Although photobleaching occurs in FingR.GFP-expressing cells over long periods, short-term imaging is possible. We were therefore able to examine puncta dynamics with FingR.GFP, and found similar puncta duration compared to GFP-tagging (Fig. 4c). Next, we examined the spatial distribution of GFP-tagged puncta by tracking them in individual OPCs (Fig. 4d). Over the course of 7 hours, GFP-Gephyrin puncta covered a large space, with a convex hull volume nearly 50 times the volume of an OPC from a single timepoint (Fig. 4d-e). Yet this convex hull volume was not significantly different from the convex hull volume of the OPC over the imaging window, suggesting that GFP-Gephyrin assembly stays tuned with OPC process remodeling (Fig. 4d-e). Puncta assembly and disassembly occurred in process regions that were relatively stable as well as those that were remodeled over the imaging window (Fig. 4f-g), suggesting that GFP-tagged puncta dynamics in OPCs were not simply caused by process remodeling. Altogether, GFP-tagged postsynaptic puncta assemble dynamically and cover a large space over time in tune with OPC process remodeling.

**Fig. 4.**
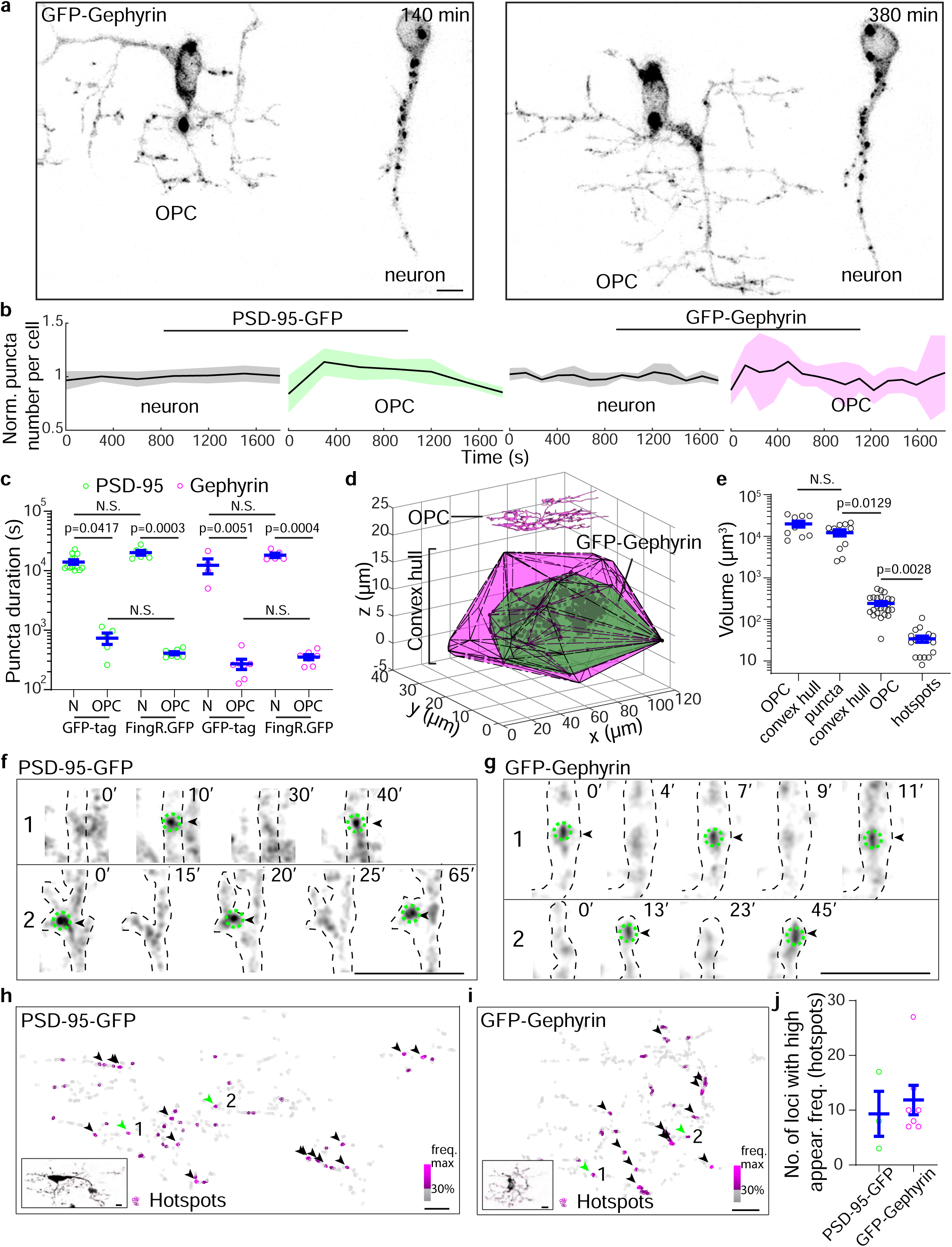
PSD-95-GFP and GFP-Gephyrin puncta in OPCs are dynamic and repeatedly assemble at hotspots. (a) Representative *in vivo* images at 2 timepoints of GFP-Gephyrin in a neuron and an OPC next to each other in the spinal cord at 3 dpf, related to Supplementary Video 1. (b) Normalized puncta number per cell of PSD-95-GFP and GFP-Gephyrin in neurons and OPCs within 30 minutes. Black line, average; shaded area, standard deviation. From left, n = 12, 3, 4, and 6 cells from 6, 10, 4, and 4 fish. (c) Puncta duration in log scale in neurons (N) and OPCs at 3 dpf for PSD-95-GFP, FingR.GFP.PSD-95, GFP-Gephyrin, and FingR.GFP.Gephyrin. n = 13, 5, 6, 7, 4, 7, 5, and 7 cells from at least 4 fish each condition. (d) Convex hull of OPC processes and GFP-Gephyrin puncta over 7 hours. The OPC process convex hull is in magenta and GFP-Gephyrin convex hull is in green. An OPC at a single timepoint is shown on top. (e) Volumes of OPC process convex hull, GFP-Gephyrin puncta convex hull, OPC at a single timepoint, and GFP-Gephyrin hotspots. From left, n= 10, 11, 23, and 18 cells from at least 10 fish. (f-g) Representative time-lapse images of (f) PSD-95-GFP and (g) GFP-Gephyrin within OPC processes. The numbers indicate the corresponding regions. #1 is in a relatively stable process; #2 is in a dynamic process over the imaging time course. The dashed lines outline the process, and arrowheads indicate puncta appearance. Time unit is in minutes. (h-i) Heatmaps of puncta appearance frequency over time for (h) PSD-95-GFP and (i) GFP-Gephyrin. 30% of max frequency is used as coloring threshold: below threshold, grey; above threshold, a shade of magenta corresponding to the frequency. Loci with high puncta appearance frequency (> 3 standard deviations above average) are marked with arrowheads. The numbers with green arrowheads indicate puncta shown in time lapse images in (f-g). Bottom left, image of the OPC with puncta marked with magenta circles. (j) Quantification of loci with high puncta appearance frequency in OPCs. From left, n = 3 and 7 cells from 3 and 5 fish. All data are represented as mean ± SEM; (c,e) Kruskal-Wallis test followed by uncorrected Dunn’s multiple comparison test; scale bars 5 μm.

Multiple appearances of puncta in the same area (Fig. 4f-g) raises the question if specific regions are preferred for synaptic assembly. To test this, we plotted PSD-95-GFP and GFP-Gephyrin puncta over time and overlaid heatmaps of puncta frequencies (Fig. 4h-i). We found approximately 10 regions (1 μm x 1 μm x 4 μm volumes) per OPC where multiple puncta appeared frequently (3 standard deviations above the average), and refer to these regions as hotspots (Fig. 4j). Hotspots represented only 0.3% of GFP-Gephyrin puncta space coverage (Fig. 4e). These results suggest that some postsynaptic puncta in OPCs assemble repeatedly and preferentially in the confined spaces of hotspots.

### GFP-Gephyrin hotspots in OPCs predict a subset of future myelin sheath formation in a synaptic release-mediated manner

In the zebrafish spinal cord, a myelinating oligodendrocyte forms an average of 10 myelin sheaths^36^, similar to the hotspot number we observed per OPC. We wondered if these hotspots in OPCs were associated with future myelin sheath formation. To test this, we conducted long-term imaging of individual OPCs that differentiated and myelinated axons (Supplementary Video 2) and measured the distances from GFP-Gephyrin hotspot centers (centroids of 1 μm x 1 μm x 4 μm) to future myelin sheaths (Fig. 5a-d, see Methods). In comparison, we selected 30 representative points outside hotspots (nonhotspot) in an OPC (See Methods). The hotspots were mostly in proximity to future myelin sheaths (Fig. 5e), and the hotspot-sheath distance was significantly smaller than the nonhotspot-sheath distance (Fig. 5f). Further, we examined the percentage of hotspots or nonhotspots that predicted where future myelin sheaths formed. We chose 1 μm as a threshold because we analyzed previous EM datasets^40^ and our IHC images and found a distance of 1.4-1.6 μm between two myelinated axons in the dorsal spinal cord (Extended Data Fig. 4a-c). Within 1 μm, 29.6 ± 8.7% of hotspots in an OPC predicted future sheath locations, while only 5.9 ± 3.4% of nonhotspots did (Fig. 5g). Consistent with this result, 31.2 ± 7.1% of puncta in the hotspot region predicted future myelin sheaths, significantly higher than the prediction by puncta outside (15.1 ± 3.4%) or by chance (5.5 ± 1.1%, myelin volume / OPC survey volume) (Extended Data Fig. 4d-e). In addition, 14.1 ± 2.7% of future myelin sheaths in oligodendrocytes were predicted by hotspots in OPCs (Extended Data Fig. 4f), suggesting that hotspots in OPCs predict a subset of myelin sheath formation.

**Fig. 5.**
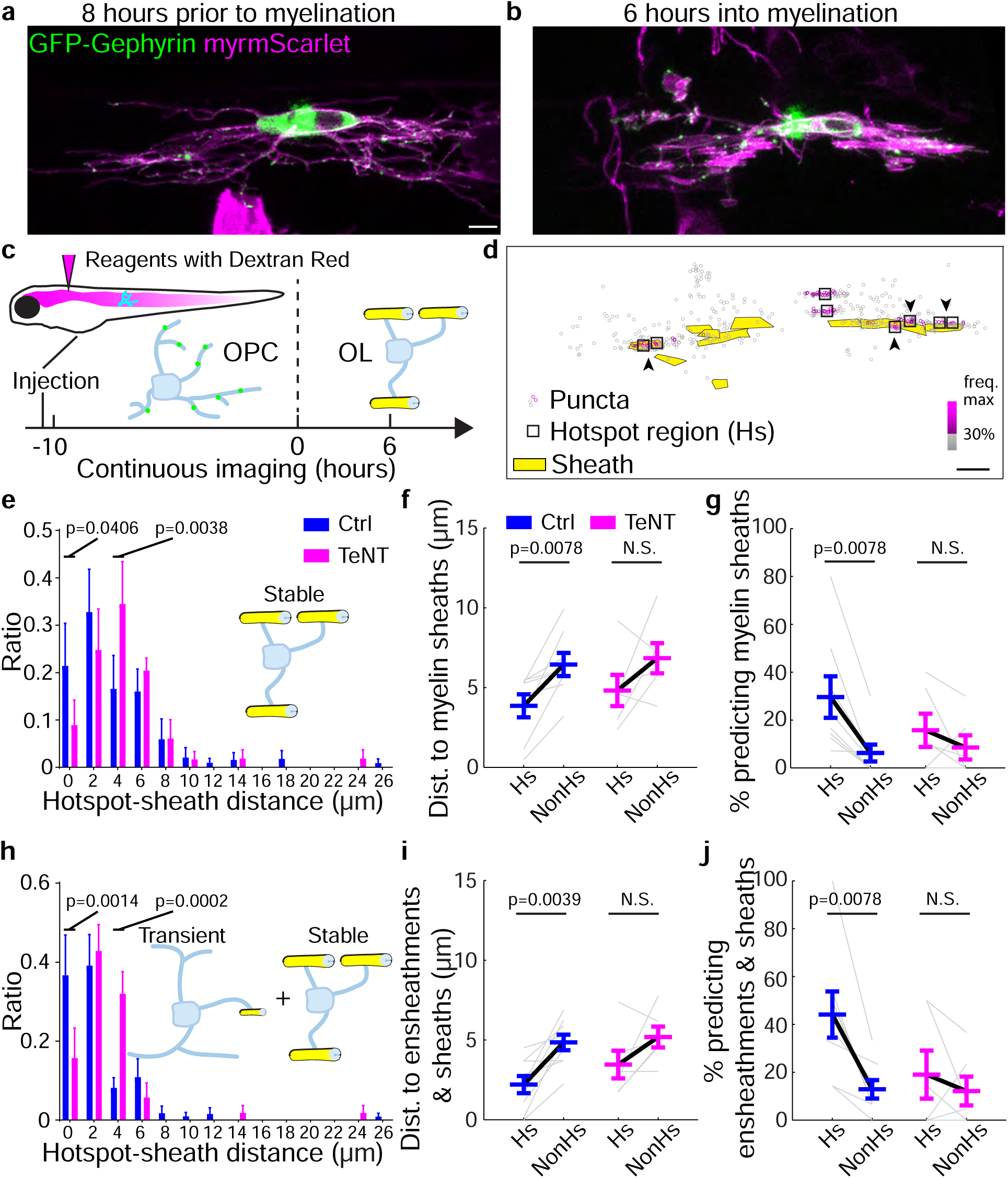
GFP-Gephyrin hotspots in OPCs predict a subset of future myelin sheath formation in a synaptic release-mediated manner. (a-b) Representative images of GFP-Gephyrin (green) in an oligodendrocyte lineage cell (magenta) at 2 different stages: (a) OPC stage that is 8 hours prior to the onset of myelination and (b) oligodendrocyte stage that is 6 hours after myelination begins, related to Supplementary Video 2. (c) Schematic showing the imaging protocol of individual OPCs that differentiate into myelinating oligodendrocytes (OLs). The first sheath appearance is designated as time 0. Images from up to 10 hours before time 0 are used to identify OPC hotspots, and images after time 0 are used to define sheath regions. TeNT is delivered through ventricle injection before imaging. (d) Single-plane merged image of GFP-Gephyrin puncta frequency heatmap at OPC stage and sheath regions at oligodendrocyte stage. Hotspot regions (black box) and sheaths (yellow) are indicated. Hotspots that exist within where myelin sheaths form are indicated with arrowheads. (e) Frequency distribution of hotspot-sheath (stable) distances with control and TeNT treatments. F[13, 168] = 11.39. (f) Quantification of hotspot (Hs)- and nonhotspot (nonHs)-sheath (stable) distances with control and TeNT treatments. control, n = 8 cells from 7 fish; TeNT, n = 6 cells from 6 fish. (g) Percentage of Hs and nonHs in an OPC that predict where myelin sheaths (stable) form within 1 μm with control and TeNT treatments. control, n = 8 cells from 7 fish; TeNT, n = 6 cells from 6 fish. (h) Frequency distribution of Hs-sheath (all) distances with control and TeNT treatments. F[13, 168] = 23.69. (i) Quantification of Hs- and nonHs-sheath (all) distances with control and TeNT treatments. control, n = 8 cells from 7 fish; TeNT, n = 6 cells from 6 fish. (j) Percentage of Hs and nonHs in an OPC that predict where myelin sheaths (all) form within 1 μm with control and TeNT treatments. control, n = 8 cells from 7 fish; TeNT, n = 6 cells from 6 fish. All data are represented as mean ± SEM; (f,g,i,j) Wilcoxon matched-pairs signed rank test; (e,h) Two-way ANOVA and Fisher’s LSD test; scale bars 5 μm.

Next, we asked if the relationship between OPC hotspots and future myelin sheaths was dependent on synaptic release. We blocked synaptic release by injecting tetanus toxin (TeNT) into ventricles between 2-3 dpf (Fig. 5c). We verified the TeNT delivery to OPCs in the CNS with Dextran injection (Extended Data Fig. 4g; see Methods). TeNT administration caused a right-shift in hotspot-sheath distance distribution (Fig. 5e). The hotspot-sheath distance was no longer significantly smaller than the nonhotspot-sheath distance (Fig. 5f). More importantly, TeNT administration reduced the hotspots’ prediction of myelin sheaths (Fig. 5g), the prediction of myelin sheath by GFP-Gephyrin puncta in the hotspot regions (Extended Data Fig. 4e), and the percentage of myelin sheaths predicted by hotspots (Extended Data Fig. 4f). This was not due to changes in OPC convex hull volume, OPC hotspot number, or myelin sheath number (Extended Data Fig. 4h-j). Together, these results suggest that GFP-Gephyrin hotspots can predict myelin sheath locations in a synapse-release dependent manner.

Recent work suggests that early myelination exhibits dynamic and transient ensheathments^41^. We similarly observed this phenomenon (Extended Data Fig. 4k) and wondered if the OPC hotspots could predict transient ensheathments. We found that within 1 μm, 32.5 ± 7.1% of hotspots in an OPC predicted future transient ensheathment locations in a synapse-release dependent manner (Extended Data Fig. 4l-n). More importantly, when we combined all ensheathments (stable sheaths and transient ensheathments), 44.2 ± 9.7% of hotspots in an OPC predicted future sheath locations in a synapse-release dependent manner (Fig. 5h-j). This percentage is higher than prediction by either stable sheaths or transient ensheathments, but lower than the combination of two, suggesting that some hotspots can predict both stable sheaths and transient ensheathments, while others predict only one type.

### OPC Ca^2+^ activity is associated with synapses and predicts myelin sheath formation

Given the role of Ca^2+^ signaling in sheath remodeling^18,42^, we examined the relationship of Ca^2+^ signaling events with synapses and sheath formation *in vivo*. With a plasma membrane-targeted GCaMP6s, we analyzed OPC Ca^2+^ activity at single-cell resolution *in vivo* (Fig. 6a-b). We found that 99% of Ca^2+^ events in OPCs occurred in MDs on processes, and 97% of OPCs exhibited only MD Ca^2+^ activity during our recordings (Fig. 6c, Supplemental Video 3-4), consistent with recent work in mammals^43,44^. We examined the proximity between Ca^2+^ events in MDs and a presynaptic marker, synaptophysin-RFP^17^. Through 3D reconstruction, we found that 42 ± 6% of MD Ca^2+^ events were within 0.5 μm proximity to synaptophysin-RFP puncta at 5 dpf (Fig. 6d-e). We also observed that some Ca^2+^ events initiated from a “puncta”-like region in proximity to synapses (Fig. 6f-g). Next, we performed time-lapse imaging to measure the distance from MD Ca^2+^ events in an OPC to sheaths formed after the OPC differentiated and myelinated axons (Fig. 6h-i). In comparison, we selected 30 representative points outside MDs (nonMD) in an OPC (See Methods). The MD-sheath distance was significantly smaller than the nonMD-sheath distance (Fig. 6j). Next, we examined the percentage of MDs that predicted where myelin sheaths formed. Within 1 μm, 32.0 ± 9.4% of MDs in an OPC predicted future sheath locations, while only 4.3 ± 1.2% of nonMDs did (Fig. 6k). These measurements of MD-future sheath are similar to those of hotspots-future sheath, suggesting that MD Ca^2+^ activity in OPCs *in vivo*, along with Gephyrin hotspots, predetermines where a subset of myelin sheaths form.

**Fig. 6.**
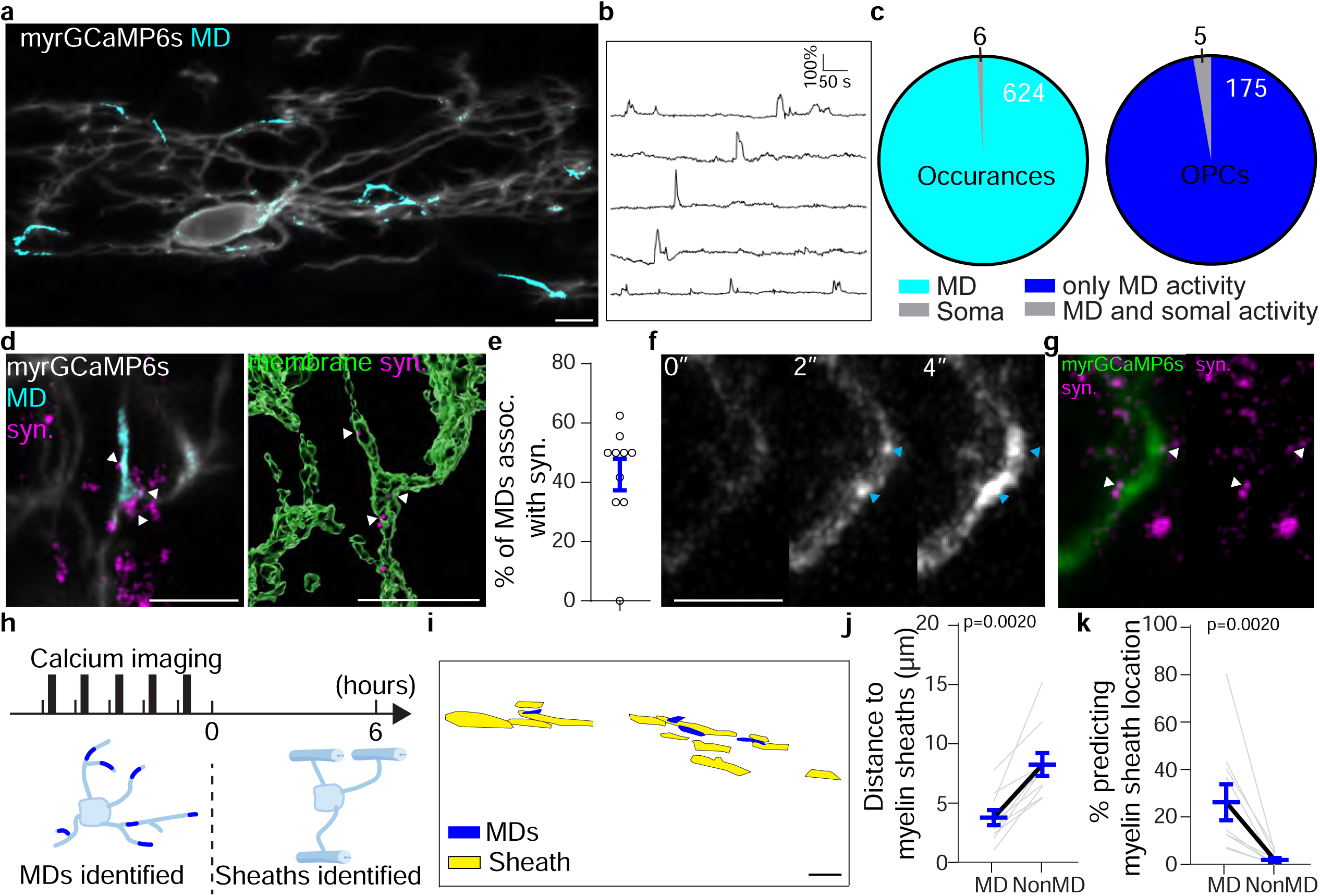
OPC Ca^2+^ activity is associated with synapses and predicts myelin sheath formation. a) Representative images of plasma membrane-targeted myrGCaMP6s in an OPC (grey, average projection) showing activity in MDs (cyan). (b) Individual traces plotting myrGCaMP6s intensity (ΔF/F_average_) in 5 MDs from (a). Scale bar, 100% and 50s. (c) Left, the occurrences of Ca^2+^ activity in MDs vs. soma; right, the number of OPCs with only MD activity vs. OPCs with both MD and somal activity. n = 180 cells from 107 fish. (d) Left, representative image of a Ca^2+^ MD (cyan) in an OPC and presynaptic synaptophysin-RFP (magenta); Right, 3-D reconstructed image showing synaptophysin-RFP puncta (magenta) that are within 0.5 μm from MDs (arrowheads). (e) The percentage of Ca^2+^ MDs that are associated with synaptophysin-RFP. n = 7 cells from 7 fish. (f) Time-lapse images of two Ca^2+^ spikes that initiate as puncta and spread to form MDs, indicated by blue arrowheads. The time is in seconds. (g) The two MDs from (f) initiate in proximity (within 0.5 μm) to presynaptic synaptophysin-RFP (magenta), indicated by white arrowheads. (h) Ca^2+^ imaging protocol of individual OPCs that differentiate into oligodendrocytes. The first sheath appearance is designated as time 0. Images up to 6 hours before time 0 are used to identify OPC Ca^2+^ events in MDs, and images around 6 hours after time 0 are used to define sheath regions. (i) Single-plane merged image of Ca^2+^ MDs at OPC stage and sheath regions at oligodendrocyte stage. Ca^2+^ MDs (blue) and sheaths (yellow) are indicated. (j) Quantification of MD- and nonMD-sheath distances. n = 10 cells from 8 fish. (k) Percentage of MDs or nonMDs in an OPC that predict where myelin sheaths form within 1 μm. n = 10 cells from 8 fish. All data are represented as mean ± SEM; (j,k) Wilcoxon matched-pairs signed rank test; scale bars 5 μm.

### Synaptic release contributes to generating OPC Ca^2+^ activity *in vivo*

Given the close relationship between Ca^2+^ activity and synapses, we investigated the role of synapses in OPC Ca^2+^ activity *in vivo.* We blocked synaptic release by sparsely expressing tetanus toxin light chain (TeNTlc)-tdtomato^45^ in axons at 5 dpf (Fig. 7a). Compared to OPCs that neighbored few TeNTlc-expressing axons (< 4, few), those that neighbored many TeNTlc-expressing axons (> 4, many) exhibited significantly reduced frequency of Ca^2+^ activity (0.013 ± 0.002 vs. 0.020 ± 0.002 Hz), peak amplitude (1.4 ± 0.08 vs. 1.81 ± 0.12 fold), basal signal intensity (0.74 ± 0.06 vs. 1.00 ± 0.10), and MD length (6.00 ± 0.25 vs. 7.03 ± 0.38 μm) (Fig. 7b-c and Extended Data Fig. 5a). Next, we delivered tetanus toxin (TeNT) proteins to the CNS through ventricle injection at 5 dpf and analyzed Ca^2+^ activity before and after injection (Fig. 7d). We coinjected Dextran (MW 3,000) and verified that Dextran surrounded OPCs before Ca^2+^ imaging (Fig. 7e). While control solution administration had no effect, TeNT administration led to nearly 45% reduction in MD event number and frequency of Ca^2+^ activity (Fig. 7f, g, Extended Data Fig. 5b and Supplementary Video 5). These results suggest an important role for synaptic release in OPC Ca^2+^ activity. TeNT administration also increased peak duration and MD area, suggesting that the synapse-dependent Ca^2+^ activity is associated with shorter duration and smaller MDs.

**Fig. 7.**
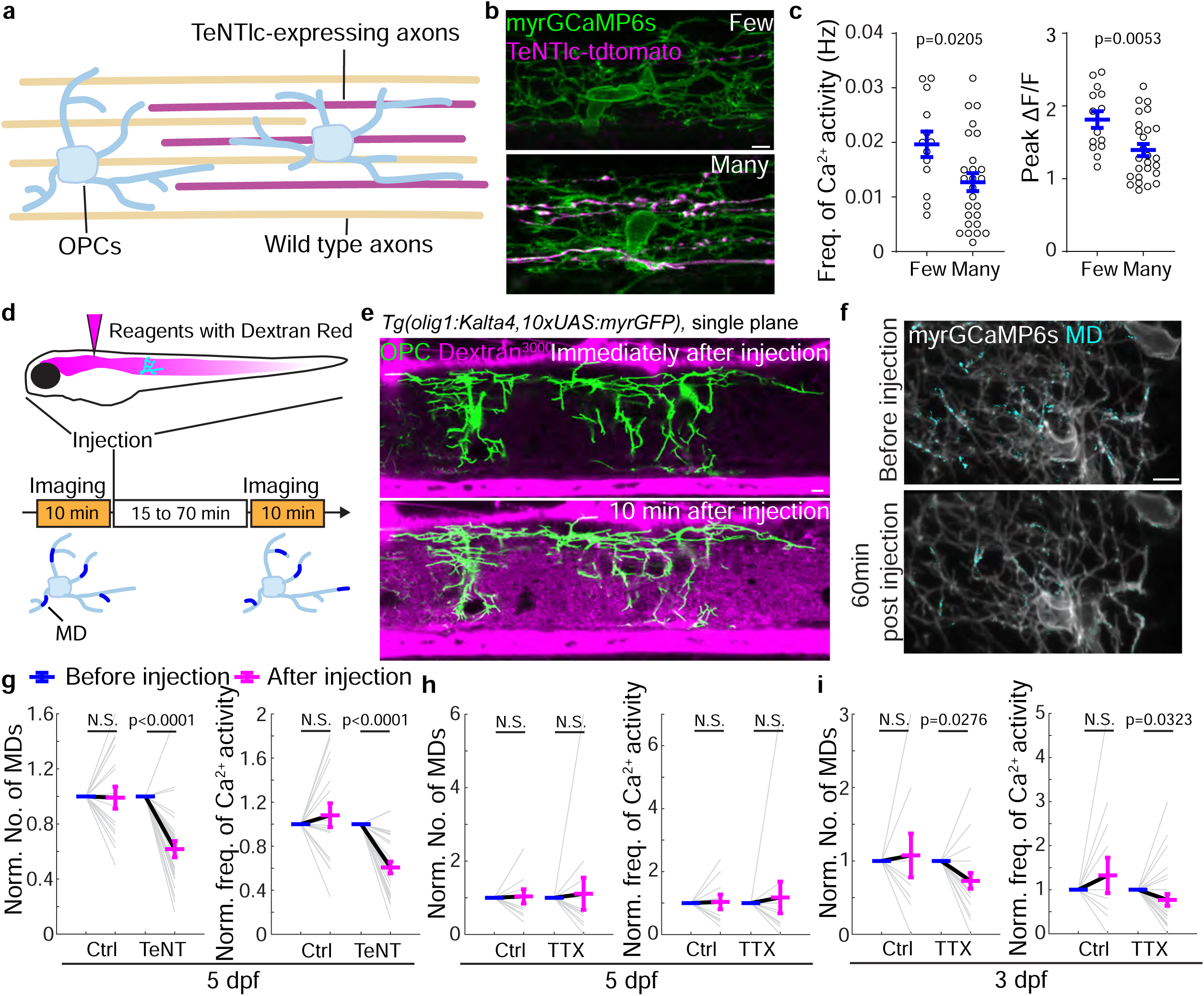
Synaptic release contributes to generating OPC Ca^2+^ activity *in vivo*. (a) Schematic showing OPCs neighboring few and many TeNTlc-expressing axons with the sparse-labeling approach. (b) Representative images of OPCs neighboring few (<4) and many (≥4) axons expressing TeNTlc-tdtomato. (c) The frequency of Ca^2+^ activity and peak amplitude in OPCs that neighbor few (<4) and many (≥4) TeNTlc-expressing axons at 5 dpf. Few, n = 14 cells from 8 fish; many, n = 26 cells from 15 fish; t[37] = 2.420 and t[38] = 2.958. (d) Experimental paradigm of reagent delivery and Ca^2+^ imaging. (e) Representative single-plane images of individual spinal cord OPCs immediately and 10 min after ventricle injection of Dextran^3000^ Texas Red (magenta) in *Tg(olig1:Kalta4,10xUAS:myrGFP)* animals. (f) Representative images of myrGCaMP6s (grey, average projection) and Ca^2+^ MDs (cyan) with control and TeNT treatment. (g) Normalized Ca^2+^ MD number and frequency of Ca^2+^ activity in OPCs before (blue) and after (magenta) injection of control or TeNT solution at 5 dpf. Paired data are indicated with grey lines. control, n = 17 from 13 fish; TeNT, n = 26 cells from 18 fish. (h) Normalized Ca^2+^ MD number and frequency of Ca^2+^ activity in OPCs before (blue) and after (magenta) injection of control or TTX solution at 5 dpf. Paired data are indicated with grey lines. From left, n = 13 and 17 cells from 6 and 9 fish. (i) Normalized Ca^2+^ MD number and frequency of Ca^2+^ activity in OPCs before (blue) and after (magenta) injection of control or TTX solution at 3 dpf. Paired data are indicated with grey lines. From left, n = 15 and 31, cells from 7 and 13 fish. All data are represented as mean ± SEM; N.S., not significant; (c) unpaired *t*-test or (g,h,i) Wilcoxon matched-pairs signed rank test; scale bars 5 μm.

These effects could be due to reduced neuronal activity and evoked synaptic release by TeNT. But to our surprise, we found that blocking neuronal action potentials with the voltage-gated Na^+^ channel blocker tetrodotoxin (TTX) had no effect on the OPC Ca^2+^ activity at 5 dpf (Fig. 7h and Extended Data Fig. 5c). We excluded the possibility of ineffective TTX delivery in three ways: we observed larval paralysis within 3 minutes post-injection (30/30), a complete block of motor neuron Ca^2+^ activity (Extended Data Fig. 5e-f), and dextran co-injected with TTX (dextran MW3000, larger than TTX) surrounded the imaged OPCs (120/120). Therefore, the reduced Ca^2+^ activity upon TeNT injection was most likely due to the effects of TeNT on spontaneous release at neuron-OPC synapses, rather than neuronal activity. We did, however, observe reduced OPC Ca^2+^ activity upon TTX injection at the earlier timepoint of 3 dpf (Fig. 7i and Extended Data Fig. 5d), consistent with a previous report^18^ and suggesting a diminishing role of neuronal activity and evoked synaptic release over development. Altogether, our results establish arole for synaptic release in generating OPC Ca^2+^ activity *in vivo*.

### Disruption of postsynaptic molecules impairs OPC development and myelination

Previous studies found that blocking neuronal activity and presynaptic release could impair myelination^10,12^. Given how GFP-Gephyrin assembly and Ca^2+^ activity in OPCs predict myelin sheath formation in a synaptic release-dependent manner, we next sought to examine the role of postsynaptic molecules in OPC development and myelination. We reasoned that disruption of these molecules could impair OPCs development and myelination because without them, OPCs would have to bypass the lack of presynaptic inputs to differentiate and choose axons for myelination. We developed and verified a cell-specific CRISPR/Cas9-mediated gene disruption system (Extended Data Fig. 6). Besides *dlg4a&b* and *gephyrinb* (See Methods), which are important for excitatory and inhibitory synapses, respectively, we also targeted *neuroligin3* (*nlgn3*), because it is highly expressed in OPCs (Extended Data Fig. 1a-b)^17,19^, localized to neuron-OPC synapses (Extended Data Fig. 7a-d), important for MAGUK and Gephyrin density (Extended Data Fig. 7e-f), and Ca^2+^ activity in OPCs (Extended Data Fig. 7g-i). To examine if OPC differentiation is affected by disruption of these postsynaptic molecules, we calculated the ratio of myelinating oligodendrocytes at different stages of development and found that *nlgn3b* disruption in oligodendrocyte lineage cells led to fewer myelinating oligodendrocytes and more OPCs (Fig. 8a-c). To assess myelination, we measured the number and total length of nascent myelin sheaths. *dlg4a*&*b-* or *gephyrinb*-disruption reduced the total length of myelin sheaths at 5 dpf (Fig. 8d-f). *nlgn3b*-disruption resulted in a reduction of the number and total length of myelin sheaths at both 3 dpf and 5 dpf (Fig. 8d-f and Extended Data Fig. 8a-b). Overexpression of a presumed dominant-negative Nlgn3b (Δ116) that lacks c-terminal domains required for interaction with MAGUK and Gephyrin^23,46^ (Extended Data Fig. 8c) also led to a reduction of the total length of myelin sheaths (Fig. 8f). In addition, among myelinating oligodendrocytes, we noted that 7.6 ± 1.8% upon *nlgn3b* disruption possessed cytoplasmic processes with abnormal trapezoidal structures (Fig. 8g and Extended Data Fig. 8d-e), suggestive of defective OPC differentiation. Together, these results support a role for postsynaptic molecules in proper OPC differentiation and myelination.

**Fig. 8.**
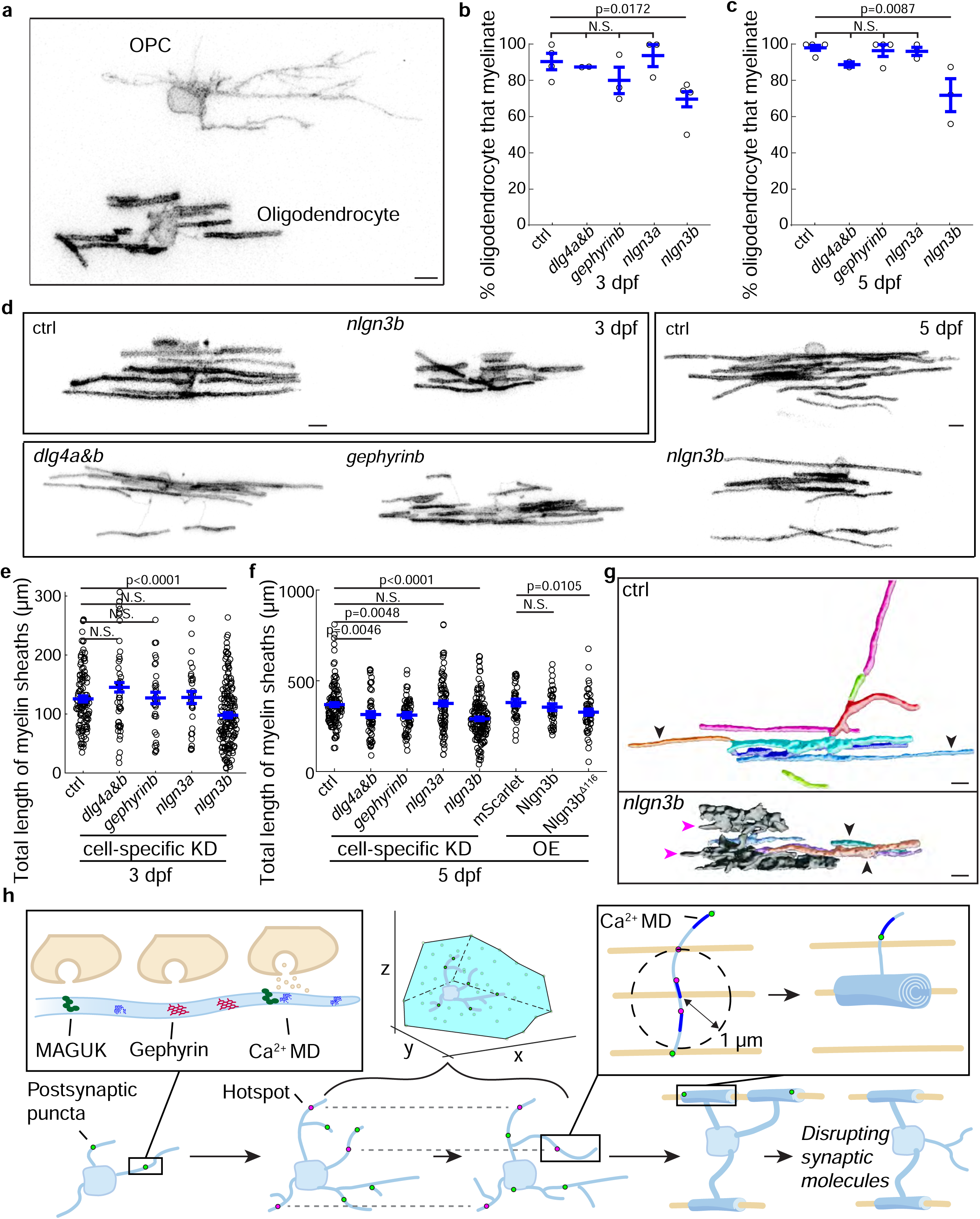
Disruption of postsynaptic molecules impairs OPC development and myelination. (a) Representative images of an OPC and an oligodendrocyte in a *nlgn3b*-targeted fish. (b-c) Percentage of oligodendrocytes that myelinate in ctrl-, *dlg4ab*-, *gephyrinb-*, *nlgn3a*-, and *nlgn3b*-targeted fish at (b) 3 dpf and (c) 5 dpf. 3 dpf, n = 144, 57, 49, 41, and 193 cells from 78, 30, 27, 26, and 97 fish; 5 dpf, n = 155, 58, 79, 84, and 198 cells from 80, 36, 40, 61, and 93 fish. (d) Representative morphologies of oligodendrocytes with control- (ctrl), *dlg4ab*-, *gephyrinb*-, and *nlgn3b*-targeted for cell-specific knockdown at 3 dpf and 5 dpf. (e) Total length of myelin sheaths per oligodendrocyte at 3 dpf in ctrl-, *dlg4ab*-, *gephyrinb-*, *nlgn3a*-, and *nlgn3b*-targeted oligodendrocytes. n = 121, 50, 34, 30, and 170 cells from 78, 30, 27, 26, and 97 fish. (f) Total length of myelin sheaths per oligodendrocyte at 5 dpf in ctrl-, *dlg4ab*-, *gephyrinb-*, *nlgn3a*-, and *nlgn3b*-targeted oligodendrocytes and mScarlet-, Nlgn3-, Nlgn3 (Δ116)-expressing oligodendrocytes. n = 118, 52, 63, 80, 152, 40, 40, and 49 cells from 80, 36, 40, 61, 93, 31, 36, and 39 fish. (g) Morphology of oligodendrocytes with ctrl- and *nlgn3b*-targeted for cell-specific knockdown at 5 dpf. Top, 3D-reconstructed single oligodendrocyte with sheath (colored) and non-sheath (gray). Sheath (black arrowheads) and processes (magenta arrowheads) are indicated. (h) *In vivo* model of neuron-OPC synapses: they are composed of MAGUK and Gephyrin and contribute to OPC Ca^2+^ activity through spontaneous release; they exhibit fast dynamics, which enable OPCs to survey large spatial volumes; synaptic assembly hotspots and associated Ca^2+^ activity at the OPC stage predict where a subset of myelin sheaths form at the oligodendrocyte stage; synaptic assembly is regulated through development while disruption of synaptic molecule*s* impairs OPC development and myelination. All data are represented as mean ± SEM; N.S., not significant; (b,c,e,f) Kruskal-Wallis test followed by uncorrected Dunn’s multiple comparisons test; scale bar 5 μm.

## DISCUSSION

The landmark discovery of neuron-OPC synapses over 20 years ago has inspired a number of studies to examine their properties with electrophysiology and EM^4^, yet we still lack basic understanding of their assembly and function. Here, we establish an *in vivo* model of these synapses in zebrafish OPCs (Fig. 8h): they are composed of postsynaptic MAGUK (PSD-95) and Gephyrin and are developmentally regulated; they exhibit a wide range of alignment with synaptic partners and dynamic synapse features; they cover large spatial volumes over time but frequently and repeatedly assemble at specific and confined “hotspots;” Gephyrin hotspots and synaptic release-mediated Ca^2+^ activity partially predict myelination patterns; and disruption of the synaptic molecules PSD-95, Gephyrin, and Nlgn3 impairs OPC development and myelination.

Previous studies suggest that neuronal activity and presynaptic release bias oligodendrocyte myelination to “active” axons^10,12^. However, it is not clear what mechanisms mediate this preference. Here, we identify a postsynaptic mechanism in OPCs that depends on presynaptic release and predicts future myelination, which might partially explain how neuronal activity regulates myelination. We observed an approximately 30% predication rate with 1 μm threshold by both GFP-Gephyrin hotspots and MD Ca^2+^ in OPCs, suggesting they may act in the same pathway to determine where myelin sheaths form. One possible scenario is that presynaptic release from neurons leads to synaptic activation in a nearby OPC process and subsequent Ca^2+^ activity in MDs. After the OPC “senses” this neuronal location, it could enhance the synaptic assembly in this location through Ca^2+^ signaling. Through such a positive-feedback mechanism, “hotspots” form, while the pre- and post-synaptic interactions influence the cytoskeleton to transform a process into a myelin sheath. 14% of stable myelin sheaths were predicted by GFP-Gephyrin hotspots, suggesting specificity in prediction and other factors or mechanisms involved, such as PSD-95 hotspots, neurotransmitters, or cell adhesion molecules. During OPC differentiation, we and others observed transient ensheathments that disappear rapidly after formation^47^. The GFP-Gephyrin hotspots predict the formation of transient ensheathments as well as the stable sheaths, suggesting that similar postsynaptic mechanisms in OPCs guide both transient ensheathment and stable myelin sheath formation. What type of presynaptic release is involved in predetermining myelination patterns? Multiple neurotransmitters and neuropeptides can regulate the development of oligodendrocyte lineage cells^5–8,48^. Therefore, different types of neurons across CNS regions may employ a combination of neurotransmitters and/or neuropeptides to influence myelination of their axons. Here, we focus on Gephyrin-containing synapses and predict that inhibitory neurotransmitters, such as glycine and GABA, may be involved in regulating the dynamic assembly of Gephyrin and OPC development. Meanwhile, how the presynaptic release sites apposed to OPC postsynaptic sites are distributed in neurons is still not fully understood. A recent study suggests presynaptic release sites are assembled in hotspots towards which myelin sheaths grow^49^. Together with our data, this suggests that both pre- and post-synaptic sites of neuron-OPC synapses are dynamically assembled at “hotspots” that predetermine where future myelin sheaths form on axons. The pre- and post-synaptic alignment at OPC synapses averages 40-50%, suggesting that some synaptic locations miss other synaptic molecules and are likely immature^50–53^. Such alignment ratios and fast synaptic assembly are in tune with an OPC’s dynamic cell biological properties and could allow these cells to effectively survey their environments.

Several studies have identified synaptic mechanisms underlying Ca^2+^ elevation in OPCs under conditions of increased neuronal activity (*e.g*., through axon stimulation) with a focus on somal activity^14–16,54^. Interestingly, recent studies provided evidence that neuronal activity or synaptic release contribute little to OPC MD Ca^2+^ activity in adult mice brains^43,44^. Like these studies, we find that OPC Ca^2+^ activity *in vivo* mainly occurs in MDs; in contrast, we show that MD Ca^2+^ activity does require synaptic release in the zebrafish spinal cord, suggesting interesting regional and/or developmental differences that will be important for future study. Neuronal activity and evoked synaptic release play a role in OPC Ca^2+^ activity at 3 dpf, but to our surprise not at 5 dpf. Considering that OPC differentiation initiates at 3 dpf, this suggests that OPCs are sensitive to neuronal activity through Ca^2+^ signaling at this critical stage of development. This sensitivity may be important for OPC differentiation and myelination in development. This development-specific sensitivity to neuronal activity also cautions the usage of axon stimulation that has been widely used in previous studies of OPC Ca^2+^ activity^14–16,54^. In our Ca^2+^ assays, we observe “outlier” OPCs that behave differently from the majority of OPCs. For example, some OPCs exhibit no change or even an increase of Ca^2+^ activity upon TeNT treatment or TTX treatment while most OPCs exhibit reduction. These “outlier” OPCs and activity could be due to Ca^2+^ generated from synapse-independent mechanisms, such as adrenergic receptor-mediated Ca^2+^ activity as recently suggested in cerebral cortex^43,44^. In line with this, nearly 61% of Ca^2+^ MDs do not associate with presynaptic markers, and nearly 55% residual Ca^2+^ activity remains after TeNT treatment, supporting synapse-independent Ca^2+^- inducing mechanisms in OPCs as well as synapse-dependent mechanisms.

Our findings that *dlg4a&b*, *gephyrinb*, or *nlgn3b* knockdown in oligodendrocyte lineage cells impairs OPC differentiation and myelination suggests a potential role of synapses in OPCs. This is consistent with our observations that GFP-Gephyrin hotspots and Ca^2+^ MDs in OPCs predict where myelin sheaths form in oligodendrocytes. The myelination defects were not detected until 5 dpf with *dlg4a&b* or *gephyrinb* knockdown, but were detected at both 3 and 5 dpf with *nlgn3b* knockdown. We reasoned that this was due to Nlgn3’s broader presence and function at Glutamatergic, GABAergic, and Glycinergic synapses. *dlg4a&b*, *gephyrinb*, or *nlgn3b* knockdown approaches likely have larger impacts on OPC activity and influence more OPCs than did disruption of single synaptic receptors from previous studies. In line with our findings, an *in vitro* study found that incubating neuron-glia co-cultures with extracellular domains of Nlgn3 inhibits OPC differentiation and axon wrapping^55^. Interestingly, disruption of the *nlgn3* homologues, *nlgn1* or *nlgn2*, in oligodendrocytes has no effects on the total myelin sheath length^23^, suggesting that synaptic disruption at the OPC stage rather than the oligodendrocyte stage affect myelination. Although non-synaptic roles of Nlgn3 have been reported in glia^56^, our results suggest a synaptic role of Nlgn3 on OPC development and myelination: close association between Nlgn3 and synapses by IHC and *in vivo* imaging; reduced MAGUK and Gephyrin puncta in OPCs upon *nlgn3b* knockdown; reduced myelination by truncated Nlgn3 that lacks MAGUK and Gephyrin interaction domains. Moreover, the similar effects on myelination from disruptions of *dlg4a&b*, *gephyrinb*, and *nlgn3b* strongly indicate a role of synapses in OPC development and myelination.

The signaling pathways downstream of synapse/Ca^2+^ MDs in OPCs remain to be elucidated. Local Ca^2+^ activity might alter the cytoskeleton via Ca^2+^-dependent kinases^57^ or affect lipid raft domains to change local signaling^54^. Notably, focal adhesion kinases (FAKs) can affect cell motility and interactions with other cells and are tightly connected to Ca^2+^ activity^58^. FAKs promotes glioma growth downstream of Nlgn3 in response to neuronal activity^3,59^, and we find that *nlgn3* disruption leads to impaired OPC differentiation, reduced oligodendrocyte myelination, and abnormal morphology. Therefore, Nlgn3 and FAKs may act downstream of synapses and Ca^2+^ signaling to regulate OPC behavior through cytoskeletal rearrangement. Overall, our *in vivo* study establishes a relationship between neuronal activity, synaptic assembly, and Ca^2+^ activity at neuron-OPC synapses, and reveals their roles in predicting where myelin sheaths form in oligodendrocytes. This model inspires future directions to investigate the pre- and post-synaptic mechanisms underlying neuron-OPC interactions that regulate OPC development and shape myelination.

## Supporting information

Video 1. Time-lapse images of GFP-Gephyrin in an OPC and neuron

Video 2. Time-lapse images of GFP-Gephyrin in a differentiating OPC

Video 3. Time-lapse images of myrGCaMP6s in an OPC

Video 4. Time-lapse images of myrGCaMP6s in an OPC with both MD and somal Ca2+ activity

Video 5. Time-lapse images of myrGCaMP6s in an OPC before and after injection of TeNT at 5 dpf

## ACKNOWLEDGMENTS

This work was supported by NMSS postdoctoral fellowship FG-1907-34613 (JL), Warren Alpert Distinguished Scholar Award (JL), and NIH/NINDS grant 1R21NS120650 (KRM). We thank Dwight Bergles, Anusha Mishra, Alex Nechiporuk, and members of the Monk lab for discussions and critical reading of the manuscript. We are indebted to Austin Forbes, Tia Perry, and Grace Halsell-Vore for fish care. We thank Rafael Almeida, Paul Brehm, Marc Freeman, Henrique von Gersdorff, Eric Gouaux, David Lyons, Lei Ma, and Teresa Nicolson for reagents and suggestions as well as Fernanda Coelho, Stefanie Kaech Petrie, and the OHSU microscopy core staff for feedback and assistance in imaging. We thank Lori Vaskalis for graphic design.

## AUTHOR CONTRIBUTIONS

J.L. conceived the project with input from K.R.M. J.L. carried out experiments and data analyses. T.M. generated constructs for *nlgn3a* and *nlgn3b* disruption, GFP-tagged Nlgn3b, and truncated Nlgn3b. T.C. provided key resources prior to their publication. J.L. and K.R.M wrote the manuscript, and all authors edited and approved of the manuscript.

## DECLARATION OF INTERESTS

The authors declare no competing interests.

## Additional information

**Supplementary Information** is available for this paper.

**Correspondence and requests for materials** should be addressed to Jiaxing Li or Kelly Monk.

## EXTENDED DATA FIGURE TITLES AND LEGENDS

**Extended Data Fig. 1. MAGUK/PSD-95, Gephyrin, and Nlgn3 are present in OPCs, related to Fig. 1**

a. RNA-seq expression of murine *Dlg4* (encodes PSD-95), *Gephyrin*, and *Nlgn3* in different CNS cell types^19^. Micro/Macro, Microglia and Macrophage; NFO, Newly-formed oligodendrocyte; MOL, Myelinating oligodendrocyte.
b. Single cell RNA-seq expression of zebrafish *dlg4* (*a*/*b*), *gephyrin* (*a*/*b*), and *nlgn3* (*a*/*b*)^17^. VLMC, Vascular lepotomeningeal cell; MOL, Myelinating oligodendrocyte.
c. Representative IHC images of PSD-95 in *Tg(olig1:Kalta4,10xUAS:myrGFP)* spinal cord at 3 dpf. From left, spinal cord sections, 3D-reconstructed regions, and two examples of puncta in OPCs at single planes. Only the postsynaptic puncta that fall within OPC process volumes are shown in 3D-reconstructed images (green spheres) and are indicated with arrowheads in examples. In each example, from left are merged channel, OPC channel and PSD-95 channel.
d. % of puncta localized to OPC process (P) and soma (S). MAGUK, n = 18 cells from 8 fish; Gephyrin, n = 40 cells from 15 fish.

All data are represented as mean ± SEM; scale bars 5 μm.

**Extended Data Fig. 2. MAGUK and Gephyrin puncta numbers in OPCs increase during early development and exhibit small differences across CNS regions, related to Fig. 1**

a. Quantification of MAGUK (green) and Gephyrin (magenta) puncta number per OPC in the spinal cord at different ages by IHC. From left, n = 20, 18, 8, 17, 24, and 8 cells from at least 6 fish per condition.
b. Quantification of MAGUK (green) and Gephyrin (magenta) puncta number per OPC at 3 dpf across different CNS regions. FB&MB, forebrain and midbrain; HB, hindbrain; SC, spinal cord. From left, n = 34, 59, 43, 34, 18, and 40 cells from at least 10 fish per condition.
c. Schematic model showing the neuron-rich (NR) and axon & synapse-rich (AR) regions and two representative images showing OPCs with soma localized to NR (top) and AR (bottom) regions in *Tg(olig1:Kalta4,10xUAS:myrGFP)* spinal cord.
d. Quantification of MAGUK (green) and Gephyrin (magenta) puncta number per OPCs in NR or AR regions at 2-3 dpf. From left, n = 14, 5, 15, and 10 cells from at least 5 fish per condition.

All data are represented as mean ± SEM; N.S., not significant; (a-b) Kruskal-Wallis test followed by Dunn’s multiple comparison test; (d) Mann-Whitney test; scale bar 5 μm.

**Extended Data Fig. 3. *In vivo* characterization of PSD-95-GFP and GFP-gephyrin, related to Fig. 3 and 4**

a. Puncta diameter of PSD-95-GFP (green) and GFP-Gephyrin (magenta) in spinal cord neurons (N) and OPCs from 2-3 dpf larvae. From left, n = 10, 11, 15, and 29 cells from at least 8 fish per condition.

(b-c) Representative *in vivo* images of (b) PSD-95-GFP or (c) GFP-Gephyrin and synaptophysin-RFP (syn.) in spinal cord OPCs at 3 dpf. From left, single plane images and an example of puncta in OPCs at single planes with 3D reconstructed regions, the merged channels, the PSD-95-GFP or GFP-Gephyrin channels, and the syn. channel. In the examples, the alignments are indicated with arrowheads.

a. (d) The % of PSD-95-GFP and GFP-gephyrin puncta in an OPC that align with presynaptic synaptophysin-RFP in the spinal cord in (b-c). n = 13 and 20 cells from at least 10 fish.

(e-f) The correlation analysis of puncta number of (e) PSD-95-GFP and (f) GFP-Gephyrin with the corresponding protein overexpression levels at 3 dpf. The linear fit extrapolated to zero overexpression level is at puncta number of 28.6 and 22.9, respectively.

Puncta number of PSD-95-GFP (green) and GFP-Gephyrin (magenta) per OPC in the spinal cord at 2 dpf and 3 dpf. From left, n = 16, 18, 29, and 14 cells from at least 5 fish per condition.
Representative IHC images of PSD-95-GFP in spinal cord OPCs at 3 dpf with GFP (green) and MAGUK (magenta) antibodies. Left, merged channels; middle and right, single channels. Arrowheads indicate the GFP puncta that colocalize with MAGUK signals; cyan arrows indicate the diffused GFP that does not colocalize with MAGUK signals.
% of GFP signal that colocalizes with MAGUK signals from (h) using Imaris. n = 4 and 6 fish per condition.

All data are represented as mean ± SEM; (a,g) Mann-Whitney test; (e,f) Pearson correlation analysis with linear regression; scale bars 5 μm.

**Extended Data Fig. 4. Gephyrin hotspots predict a subset of myelin sheath formation in a synaptic release-mediated manner, related to Fig. 5**

a. Representative EM image of zebrafish dorsal spinal cord transverse section at 5 dpf^40^. Myelinated axons are indicated with magenta dots.
b. Representative IHC images of zebrafish dorsal spinal cord transverse section at 5 dpf in *Tg(mbp:GFP-caax)*. Myelin sheaths that wrap axons form circle structures (“sheath”).
c. Quantification of the nearest distance between dorsal myelinated axons from (a) and (b). n = 16 and 9 sections from 1 and 6 fish; t[23]=1.627.
d. Schematic showing the puncta inside hotspots (PIH) and the puncta outside hotspots (POH).
e. Percentage of GFP-Gephyrin PIH and POH that predict future myelin sheaths within 1 μm and percentage of stable myelin sheath volume over the entire volume of OPC with control and TeNT treatments.
f. Percentage of future stable myelin sheaths in oligodendrocytes that are predicted by hotspots in OPCs (within 1 μm) with control and TeNT treatments. n = 6 cells from 6 fish.
g. Representative image of individual spinal cord OPCs 50 min after ventricle injection of Dextran^150,000^ Antonia Red (magenta) in *Tg(olig1:Kalta4,10xUAS:myrGcamp6s)* fish.

(h-j) Quantification of (h) convex hull volume, (i) hotspot number, and (j) sheath number in control and TeNT treatment conditions. (h) n = 11 and 6 cells from 10 and 6 fish; (i) n = 18 and 6 cells from 16 and 6 fish; (j) n = 11 and 6 cells from 10 and 6 fish.

Representative time-lapse images of transient ensheathments after time 0 in *Tg(sox10:Kalta4,10xUAS:myrmScarlet)*. Arrowheads indicate the sheath appearances. Time unit is in minutes.
Frequency distribution of hotspot-transient ensheathment distances with control and TeNT treatments. F[14, 180] = 23.87.
Quantification of Hs- and nonHs-sheath (stable) distances with control and TeNT treatments. control, n = 8 cells from 7 fish; TeNT, n = 6 cells from 6 fish.
Percentage of Hs or nonHs in an OPC that predict where transient ensheathments form within 1 μm with control and TeNT treatments. control, n = 8 cells from 7 fish; TeNT n = 6 cells from 6 fish.

All data are represented as mean ± SEM; (c) unpaired *t*-test; (e) Friedman test with Dunn’s test; (f,h,i,j) Mann-Whitney test; (l) Two-way ANOVA and Fisher’s LSD test; (m,n) Wilcoxon matched-pairs signed rank test; scale bars 5 μm.

**Extended Data Fig. 5. The role of synaptic release and neuronal activity in generating OPC Ca^2+^ activity, related to Fig. 7**

a. Basal GCaMP6s intensity and MD feret’s diameter in OPCs that neighbor few (< 4) and many (≥ 4) TeNTlc-expressing axons at 5 dpf. Few, n = 14 cells from 8 fish; many, n= 26 cells from 15 fish; t[38] =

2.375 and t[38] = 2.351.

Normalized peak amplitude, peak duration, area of Ca^2+^ MD, basal intensity, and MD Feret’s diameter in OPCs before (blue) and after (magenta) injection of control or TeNT solution at 5 dpf. Paired data are indicated with grey lines. Control, n = 17 cells from 13 fish; TeNT, n = 26 cells from 18 fish.

(c-d) Normalized peak amplitude, peak duration, and area of Ca^2+^ MD in OPCs before (blue) and after (magenta) injection of control or TTX solution at (c) 5 dpf and (d) 3 dpf. Paired data are indicated with grey lines. From left, n = 15, 31, 13, and 17 cells from 7, 13, 6, and 9 fish.

Representative images of a primary motor neuron visualized by injecting *mnx1:gal4* into *Tg(10xUAS:myrGCaMP6s)*.
Normalized frequency of Ca^2+^ activity in motor neurons before (blue) and after (magenta) injection of control or TTX solution at 5 dpf. n = 7 and 6 cells from 5 and 5 fish.

All data are represented as mean ± SEM; N.S., not significant; (a) unpaired *t*-test or (b,c,d,f) Wilcoxon matched-pairs signed rank test; scale bars 5 μm.

**Extended Data Fig. 6. Cell-specific knockdown system is efficient to disrupt genes in oligodendrocytes in zebrafish, related to Fig. 8**

a. Schematic of the plasmid (10xUAS:myrmScarlet-p2A-Cas9, U6:sgRNA1;U6:sgRNA2) used to induce cell-specific Cas9 expression with membrane labeling and 2 separate U6-driven sgRNAs.
b. Representative gel images of digested PCR product of genomic regions targeted by the sgRNAs. Each lane represents a single embryo at 1 dpf. The left 4 lanes are uninjected (-) and the right 4 lanes are injected (+) with sgRNAs and Cas9 protein.
c. Schematic of larva with sparsely labeled oligodendrocyte lineage cells resulting from *Tg(sox10:Kalta4)* crossed with transgenic fish carrying the plasmid from (a).
d. Representative images of single oligodendrocytes in the spinal cord of *Tg(10xUAS:myrmScarlet-p2A-Cas9, U6:ctrl-sgRNA1;U6:ctrl-sgRNA2)* and *Tg(10xUAS:myrmScarlet-p2A-Cas9, U6:myrf^ex^*^3^*-sgRNA;U6:myrf^ex^*^4^*-sgRNA)* at 6 dpf. Knockdown of Myrf serves as a control for our targeting approach.
e. Average myelin sheath length in (d). n = 24 and 22 cells from 15 fish; t[44] = 2.247.

All data are represented as mean ± SEM; N.S., not significant; (e) unpaired *t*-test; scale bars 5 μm.

**Extended Data Fig. 7. Nlgn3 is localized to synapses and knock-down reduces postsynaptic puncta in OPCs, related to Fig. 8**

a. Representative single-plane IHC images of Nlgn3 (Af1010^60^) in OPCs with presynaptic marker SV2 in *Tg(olig1:myrmScarlet)* spinal cord at 5 dpf. From left, spinal cord sections and two alignment examples with SV2. In each example, from left are 3D reconstructed regions, the merged channels, the OPC channel, the Nlgn3 channel, and the SV2 channel. Only the postsynaptic puncta that fall within OPC process volumes are shown in 3D-reconstructed images (green spheres) with aligned SV2 (magenta spheres) and are indicated with arrowheads in other channels.
b. The number of Nlgn3 puncta in an OPC that align with presynaptic SV2 in the spinal cord at 5 dpf from control and F0 larvae with *nlgn3a and nlgn3b* knocked down by sgRNA targeting *nlgn3a^ex^*^4^ and *nlgn3b^ex^*^1^ and Cas9 protein. From left, n= 6 and 7 cells from at least 5 fish each condition.
c. Representative *in vivo* images of Nlgn3b-GFP and synaptophysin-RFP (syn.) in the spinal cord of a *Tg(olig1:Kalta4)* larva at 5 dpf. From left, single plane image and two examples of puncta in OPCs at single planes with 3D reconstructed regions, the merged channels, the Nlgn3b-GFP channel, and the syn. channel. In the examples, the aligned Nlgn3b puncta are indicated with arrowheads.
d. Number of Nlgn3b-GFP puncta in OPCs that align with synaptophysin-RFP puncta. n=6 cells from 6 fish.
e. Representative IHC images of MAGUK and Gephyrin in *Tg(sox10:Kalta4,10xUAS:myrmScarlet-p2A-Cas9)* spinal cord at 3 dpf with ctrl- and *nlgn3b*-targeted for cell-specific knockdown. From left, spinal cord sections and 3D-reconstructed images of enlarged regions. In 3D-reconstructed images, only the postsynaptic puncta that fall within OPC process volume are shown (green sphere).
f. Density of MAGUK and Gephyrin puncta in OPCs from (e). From left, n = 12, 12, 16, and 16 cells from at least 8 fish.
g. Representative *in vivo* images of an OPC expressing the cell-specific knockdown construct in *Tg(olig1:Kalta4, olig1:myrGCaMP6s)* at 4 dpf.

(h-i) The frequency of (h) Ca^2+^ activity and (i) MD number in ctrl- and *nlgn3b*-targeted OPCs at 3-4 dpf. ctrl, n = 13 cells from 13 fish; *nlgn3b*, n = 17 cells from 16 fish; t[28] = 2.260 and t[28] = 2.511.

All data are represented as mean ± SEM; (b,f) Mann-Whitney test; (h,i) unpaired *t*-test; scale bars 5 μm.

**Extended Data Fig. 8. Disruption of *nlgn3b* impairs OPC development and myelination, related to Fig. 8**

(a-b) Number of myelin sheaths per oligodendrocyte in ctrl-, *dlg4ab*-, *gephyrinb-*, *nlgn3a*-, and *nlgn3b*-targeted oligodendrocytes at (a) 3 dpf and (b) 5dpf. a, n = 121, 50, 34, 30, and 170 cells from 78, 30, 27, 26, and 97 fish; b, n = 118, 52, 63, 80, and 152 cells from 80, 36, 40, 61, and 93 fish.

Schematic showing the domains of Nlgn3b and the region mutated in the dominant-negative construct employed in our studies.
4 single planes of a trapezoid-shaped process in an oligodendrocyte in *nlgn3b*-targeted larvae at 5 dpf. Myelin sheaths are indicated by black arrowheads and abnormal processes by magenta arrowheads.
Percentage of oligodendrocytes that possess both myelin sheath(s) and abnormal processes in ctrl-, *nlgn3a*-, and *nlgn3b*-targeted fish at 5 dpf. n = 119, 84, and 170 cells from 80, 61, and 93 fish, F[6] = 13.52.

All data are represented as mean ± SEM; N.S., not significant; (a,b,e) Kruskal-Wallis test followed by uncorrected Dunn’s multiple comparisons test; scale bars 5 μm.

## SUPPLEMENTARY FIGURES WITH TITLES AND LEGENDS

**Supplementary Fig. 1. Antibody verification in zebrafish for Gephyrin (SS147011) and Nlgn3b (Af1010).**

a. IHC images of Gephyrin in the 5 dpf spinal cord in control (ctrl) and F0 larvae with sgRNA targeting *gephyrina^ex^*^2^, *gephyrinb^ex^*^7^, or *gephyrina^ex2^*&*gephyrinb^ex7^* plus Cas9 protein.
b. Quantification of Gephyrin intensity with ctrl and sgRNA targeting *gephyrina^ex2^*&*gephyrinb^ex7^* from (a). n = 5 and 9 fish for control and knockdown, respectively; t[12] = 7.322.
c. Representative transverse spinal cord IHC images of Nlgn3 at 5 dpf in wild-type and F0 larvae with *nlgn3a and nlgn3b* knocked down by sgRNA targeting *nlgn3a^ex4^* and *nlgn3b^ex1^* and Cas9 protein.
d. Quantification of Nlgn3 intensity in OPCs from (c). From left, n= 6 and 7 cells from at least 5 fish each condition.

All data are represented as mean ± SEM; Mann-Whitney test; scale bars 5 μm.

## SUPPLEMENTARY VIDEOS WITH TITLES AND LEGENDS

**Video 1. Time-lapse images of GFP-Gephyrin in an OPC and neuron**

Representative video of 10xUAS:GFP-Gephyrin in an OPC and a neuron next to each other in the spinal cord of a *Tg(sox10:Kalta4)* larva at 3 dpf. The video was generated through maximum projection of stacks.

**Video 2. Time-lapse images of GFP-Gephyrin in a differentiating OPC**

Representative video of 10xUAS:GFP-Gephyrin in an OPC that eventually differentiates in the spinal cord of a *Tg(sox10:Kalta4, 10xUAS:myrmScarlet)* larva at 3 dpf. The video was generated through maximum projection of stacks.

**Video 3. Time-lapse images of myrGCaMP6s in an OPC**

Representative video of Ca^2+^ activity in an OPC in a *Tg(olig1:Kalta4,10xUAS:myrGCaMP6s)* larva at 5 dpf.

**Video 4. Time-lapse images of myrGCaMP6s in an OPC with both MD and somal Ca^2+^ activity** Video of Ca^2+^ activity in an OPC with multiple events including a somal event in a *Tg(olig1:Kalta4,10xUAS:myrGCaMP6s)* larva at 5 dpf.

**Video 5. Time-lapse images of myrGCaMP6s in an OPC before and after injection of TeNT at 5 dpf** Representative 10-minute video of Ca^2+^ activity in an OPC before (left) and after (right) injection of TeNT in a *Tg(olig1:Kalta4,10xUAS:myrGCaMP6s)* larva at 5 dpf.

## SUPPLEMENTARY TABLES WITH TITLES AND LEGENDS

**Table 1. A list of primers used in this study**

## METHODS

### RESOURCE AVAILABILITY

#### Lead contact

Further information and requests for resources and reagents should be directed to and will be fulfilled by the lead contacts, Jiaxing Li or Kelly Monk.

#### Materials availability

All reagents generated in this paper will be shared by the lead contacts upon request. Data and code availability

All data reported in this paper will be shared by the lead contacts upon request. All original code is available from the lead contacts upon request.

Any additional information required to reanalyze the data reported in this paper is available from the lead contacts upon request.

### EXPERIMENTAL MODEL AND SUBJECT DETAILS

We used the following existing zebrafish lines and strains: *Tg(sox10:Kalta4)*^61^, *Tg(olig1:Kalta4)*^17^, *Tg(huc:synaptophysin-RFP)*^17^, *Tg(olig2:dsRed)*, *Tg(sox10:GFP-caax)*, *nacre*, and AB. The following lines were newly generated for this study: *Tg(10xUAS:myrGCaMP6s)*, *Tg(olig1:myrGCaMP6s), Tg(10xUAS:myrGFP), Tg(10xUAS:myrmScarlet), Tg(olig1:myrmScarlet)*. All experimental fish were crossed into the *nacre* background for optimal optics without pigmentation. The Kalta4/UAS system allowed sparse labeling and single cell analysis. The genotypes of fish used for each experiment are indicated in figure legends. Animals in the fish facility were maintained at 28° C with a 141’h/101’h light/dark cycle. Larvae and juveniles were nurtured with rotifer suspension and dry food (Gemma 75 and 150, respectively). Adults were fed with a combination of brine shrimp and dry food (Gemma 300). Male and female fish of 3 to 12 months-old were mated; larval fish of 2 dpf to 5 dpf were used for experiments (sex is not determined at these stages in zebrafish). All zebrafish experiments and procedures were performed in compliance with institutional ethical regulations for animal testing and research at Oregon Health and Science University (OHSU). Experiments were approved by the Institutional Animal Care and Use Committee of OHSU. Animal and cell numbers are indicated in figure legends.

### METHOD DETAILS

#### Tissue sectioning and immunohistochemistry

To prepare transverse sections, zebrafish larvae at indicated stages were first anesthetized with tricaine and then fixed in 4% PFA, 0.1% Triton at 4° C overnight. Heads and tails were removed to enable better penetration of fixative. The fish were then washed in PBS, 0.1% Triton for 5 minutes and passed in 10%, 20%, and 30% sucrose in PBS, followed by OCT embedding and cryosectioning of 20-25 μm slices. The sections were brought to room temperature for 45 minutes and rehydrated in PBS, 0.2% Triton for 1 hour. 3% BSA, 5% NGS, 0.2% Triton in PBS was used as blocking buffer. Primary antibodies: chicken anti-GFP (Invitrogen A10262), 1:1000, mouse anti-Pan-MAGUK (NeuroMab K28/86) 1:500, mouse anti-Gephyrin (Synaptic Systems 147011) 1:1000, rabbit anti-Synapsin 1/2 (Synaptic Systems 106002) 1:1000, mouse anti PSD-95 6G6-1C9 (MAB1596) 1:1000, rabbit anti-Glycine receptor (Synaptic Systems 146008) 1:1000, mouse, anti-SV2 (DSHB AB_2315387) 1:1000, rabbit anti-Nlgn3 (nittobo medical Af1010) 1:1000.

The MAGUK antibody was previously validated in zebrafish^62–64^, and we validated Gephyrin and Nlgn3 antibody in zebrafish (Supplementary Fig. 1). Secondary antibodies: A488 anti-chicken, A594 anti-rabbit, A647 anti-mouse IgGI, A647 anti-mouse (Thermo Fisher). The stained samples were mounted in prolong Glass (Invitrogen P36980) for at least a day before imaging.

#### Plasmid construction

In zebrafish, *gephyrin* is duplicated (*gephyrina* and *gephyrinb*), and there are multiple isoforms for both genes. All isoforms are almost identical in protein sequence except for the inclusion or exclusion of C3 and C4 cassettes. We amplified several isoforms from cDNA and decided to follow up two: *gephyrinb* isoform X6 (XM_021468341) because it encodes a protein that is 96% identical and 98% similar to the rat P1 isoform (NP_074056.2)^65^ that has been used to track synapses previously; and, *gephyrina* isoform X11 (XM_017351650.2) because it contains a C3 cassette previously reported to be present in glia cells^66^.

Total RNA was extracted from 3 dpf zebrafish AB larvae (50 fish) using RNAeasy Plus Kit (Qiagen 74134). cDNA was synthesized using SuperScript IV Reverse Transcriptase (Invitrogen). Two sets of primers (Supplementary Table 1) were used to clone the full length*gephyrina* and *gephyrinb* using high fidelity Q5 DNA polymerase, followed by A-addition and TA cloning into pCRII-TOPO backbones. Using Gibson assembly (NEB E5510S), we added EGFP to the 5’ end and a linker GGGGSGGGGSGGGGS in between, and integrated it into pCS 2.1 backbones with an AVRII digestion site. The 5’ end was chosen based on previous imaging studies^65^. In transient injections, GFP-Gephyrina and GFP-Gephyrinb exhibited similar expression profiles; therefore, we used GFP-Gephyrinb for our studies and refer to the GFP-3xGGGGS-Gephyrinb as GFP-Gephyrin throughout the manuscript. Using Gateway BP reaction (Invitrogen 11786100), we generated pME_ATG-GFP-Gephyrinb. Using AVRII and KPNI-HF, we cut GFP-Gephyrinb (without an ATG) from the pCS 2.1 backbone and inserted it in frame into p3E_p2A to generate p3E_p2A-GFP-Gephyrinb.

*PSD-95*/*dlg4b* transcript NM_214728.1 encodes a protein that is 76% identical and 83% similar to rat *Dlg4* (NP_062567.1) and was thus chosen for use in our studies. We added GFP to the 3’end because N-terminal palmitoylation of PSD-95 is necessary for its synaptic localization, and receptor clustering^67,68^. It was cloned from cDNA with primers (Supplementary Table 1) similar to *gephyrin* as described above. We refer to PSD-95-3xGGGGS-GFP as PSD-95-GFP. We generated pME_ATG-PSD-95-GFP and p3E_p2A-PSD-95-GFP constructs.

*nlgn3b* transcript XM_005165182 encodes a protein that is 72% identical and 81% similar to mouse *Nlgn3* (NP_766520). We synthesized the coding region from XM_005165182 using IDT Gene Synthesis. For GFP-labeling, we inserted GFP at 3’ end with a linker and generated pME_Nlgn3-GFP. In addition, we generated pME_Nlgn3 (full length) and pME_Nlgn3 (Δ116) that misses both domains for interactions with MAGUK and Gephyrin^23,46^.

pCAG_FingR.GFP-Gephyrin-CCR5TC (Addgene 46296) and pCAG_FingR.GFP-PSD-95-CCR5TC (Addgene 46295) were used to amplify the coding region with additional 5’ zinc finger binding sequence to regulate their expression^20^, followed by BP reaction to generate pME_zincfinger_FingR.GFP-PSD-95-CCR5TC and pME_zincfinger_FingR.GFP-Gephyrin-CCR5TC constructs.

pME_myrGFP, pME_myrmScarlet and pME_myrRFP-T were generated using PCR and BP reaction, using Fyn myristoylation and palmitoylation domain. p3E_mScarletcaax was generated using PCR and BP reaction. pGP-CMV-GCaMP6s^69^ (Addgene 40753) was used as a template and 3 primers were used in a Q5 PCR (15 cycles at 71° C and 20 cycles at 72° C) to amplify GCaMP6s and add a Kozak sequence, myr (Fyn myristoylation and palmitoylation domain), attB1, and attB2 sequences. This PCR product was used to generate pME_myrGCaMP6s using BP reaction.

We obtained multiple plasmids from Tol2 kits: pDestTol2pA2, pDONR221,p5E_10xUAS, p3E_polyA, p3E_p2A^70^, p5E_sox10_7.2kb_^61^, pDestTol2pACryGFP (pDestTol2GE), pDestTol2pACrymCherry (pDestTol2RE).

Using Gateway LR reactions (Invitrogen 11791020), we generated the following constructs: GE-10xUAS:myrEGFP, RE-10xUAS:myrmScarlet, PA2-olig1:myrmScarlet, GE-10xUAS:myrGCaMP6s, PA2-olig1:myrGCaMP6s, GE-10xUAS:GFP-Gephyrinb, RE-10xUAS:GFP-Gephyrina, GE-10xUAS:PSD-95-GFP, PA2-sox10:myrRFP-T-P2A-GFP-Gephyrinb, PA2-sox10:myrRFP-T-P2A-PSD95-GFP, GE-10xUAS:zincfinger_FingR.GFP-PSD-95-CCR5TC, GE-10xUAS:zincfinger_FingR.GFP-Gephyrin-CCR5TC, GE-10xUAS:Nlgn3b-GFP, RE-10xUAS:Nlgn3-P2A-mScarletcaax, RE-10xUAS:Nlgn3(Δ116)-P2A-mScarletcaax. We used Mnx1:Gal4^71^ to drive myrGCaMP6s expression in motor neurons. We used Huc:Kalta4^61^, UAS:TeNTlc-tdtomato^45^ to drive TeNTlc expression in neurons. All constructs generated in this study were fully sequenced (except *sox10* and *olig1* promoters).

#### Cell specific CRISPR/Cas9 plasmid construction and CRISPR/Cas9-mediated gene disruption

We used 5xUAS:hsp70promoter-Cas9-T2A-GFP, U6:sgRNA1;U6:sgRNA2^72^ (Addgene 74009) as a backbone and performed the following modifications: replacement of the 5xUAS:hsp70 promoter with the 10xUAS:E1b-bactin-Kozak promoter because we found the original promoter expressed in many non-specific cell types when injected into *Tg(sox10:Kalta4)*; replacement of T2A-GFP with myrmScarlet-p2A and added to the 5’end of Cas9; addition of polyA to the 3’ end of Cas9. Four sets of primers and 2 previously made plasmids (Tol2-10xUAS:E1b-mScarlet and Tol2-10xUAS:cas9-P2A-RFPT-caax) were used to generate 4 segments. These segments were integrated into the backbone by NEBuilder HiFi assembly (NEB E2621S). The new construct is referred to as 10xUAS:myrmScarlet-p2A-Cas9, U6:sgRNA1;U6:sgRNA2.

CHOPCHOP and Crisprscan were used to identify sgRNA sequences for each gene of interest: *dlg4a* exon 6, GGTGACCCACAGTCAGGCGGTGG; *dlg4b* exon 8, GGAAGCAGGACCGATCGTCCGG; *gephyrina* exon 2, GGTCATGAACGAGATCTTTGAGG; *gephyrinb* exon 7, GGAGAGCCGGCAGGATGAACTGG; *gephyrinb* exon 8, GGGCGTCCAGGTTCTTCCGCGGG; *myrf* exon 3, GACCCAACGGCGGTGCACCCGGG; *myrf* exon 4, GCTGGAGTCCGGAGGAGACTCGG; *nlgn3a* exon 1, GGTGTATGATGCTTGTCCGGGGG; *nlgn3a* exon 4, GCCTTGTGTACTTAACCGGCTGG; *nlgn3b* exon 1, GATGCGGGTCGCTGTTGCCACGG; *nlgn3b* exon 3, GGTAGTGTGCTGGCCAGCTACGG. The cutting efficiencies of these sgRNAs were verified (>95%) via enzyme digestion of the targeted locus PCR product (CRISPRtest primers, Extended Data Fig. 6b). F0 larvae with *gephyrina* and *gephyrinb* sgRNA injected together with 1 ng Cas9 protein were used to examine the specificity of the Gephyrin antibody. *gephyrinb* disruption was sufficient to eliminate Gephyrin protein in the CNS (Supplementary Fig. 1a-b). BsaI and BsmBI were used to insert the sgRNA into 10xUAS:myrmScarlet-p2A-Cas9, U6:sgRNA1;U6:sgRNA2, while the uninserted plasmid was used as controls. The offspringlarvae with sparsely labeled oligodendrocyte were examined for morphology and sheath characteristics.

#### Sparse labeling and generation of transgenic lines

Fertilized zebrafish eggs at the 1-cell stage were microinjected with 11’nL of an injection solution containing 201’ng/µL plasmid DNA, 251’ng/µL Tol2 transposase mRNA, 0.02% phenol red and 0.2M KCl. Injected F0 animals were either used for single-cell analysis or raised to adulthood to generate transgenic lines. To generate stable lines, F1 progenies were screened for germline transmission of the fluorescent reporters. F1 animals were further outcrossed and maintained in the *nacre* background to establish stable transgenic lines with single copy insertion.

#### Mounting larvae for *in vivo* imaging

For most live imaging experiments, larvae were anesthetized with 0.16 mg/mL (600μM) tricaine in embryo medium. The fish were mounted in 0.8% low melting agarose (Sigma A9414) on a coverslip for morphology and sheath examination or in 1.5% low melting agarose in a dish for time lapse imaging (hours). For Gephyrin hotspot and myrGCaMP6s imaging, larvae were immobilized with 0.5 mg/ml non-depolarizing acetylcholine receptor antagonist mivacurium chloride (Abcam) and mounted in 1.5% low melting agarose in a dish.

#### Confocal microscopy

All imaging was performed on an upright Zeiss LSM 980 confocal with Airyscan 2 in 4Y fast mode. For fixed tissues, a 63x/1.4 NA oil objective was used, and for live imaging, a 20x/1.0 NA water objective was used. All images were taken in Zeiss software. We used the following lasers: 488 nm for EGFP, myrGCaMP6s, and Alexa488; 561 nm for RFP, mScarlet, tdtomato, and Alexa594; 639 nm for Alexa647. Fixed tissue was imaged with 0.15 μm z spacing with 1.6x to 2.5x zoom. For *in vivo* analyses, PSD-95-GFP and GFP-Gephyrin were imaged with 2 μm z-spacing at an interval of 5 min (for 30 min to 1 hour) or 20 min (for longer than 1 hour) at a zoom between 3.7x-10x depending on OPC cell size. myrGCaMP6s was imaged with 2 μm z-spacing at 0.5 Hz (4 planes scanned) for 10 min or 20 min (drug injection experiments) as previously described^17,18^. Images acquired at 0.5 Hz were analyzed along with images acquired at 10 Hz with a single plane. No major differences were found on the Ca^2+^ spike characteristics, but 0.5 Hz acquisition allowed inclusion of more stacks and increased the number of events detected. Therefore, 0.5 Hz was used throughout the study. For tracking OPCs that differentiate into myelinating oligodendrocyte, Ca^2+^ imaging was done repeatedly for 5 min at the beginning of each hour.

#### Drug delivery through ventricle injection

For myrGCaMP6s experiments involving drug delivery, we injected drugs via ventricle injection as described previously^18^. In TTX treatment, following 10 min GCaMP6s imaging, a 1 nL mixture containing 0.2M KCL, 0.5% Dextran Texas Red 3000 MW (Thermo), and H_2_O (control) or TTX (0.5 mM, kind gift from Henrique von Gersdorff’s lab) were pressure injected into the hindbrain ventricle using an ASI MPPI-3. Between 15 and 35 min after injection, Dextran was imaged, followed by another 10 min of GCaMP6s imaging. Dextran surrounding the imaged OPCs was used to verify the drug delivery. TTX caused fish paralysis, further verifying efficient delivery of active drugs. After imaging, fish were removed from mounting media. Larvae injected with control solution recovered full swimming activity within 3 minutes, and TTX-injected paralyzed larvae maintained normal heartbeats for at least 3 hours. In TeNT treatment, an 8 nL mixture containing 0.2M KCL, 0.5% Dextran Antonia Red 150,000 MW (Sigma), and TeNT (listlabs 190A, 0.5 μM) were pressure injected into the hindbrain ventricle. Between 50 and 70 min after injection, Dextran was imaged, followed by another 10 min of GCaMP6s imaging. TeNT-injected larvae showed reduced swimming activity.

### QUANTIFICATION AND STATISTICAL ANALYSIS

#### OPC labeling during development

We used *sox10* and *olig1* to label OPCs. *sox10* drives expression in both OPCs and myelinating oligodendrocytes while *olig1* labels only OPCs. OPCs and myelinating oligodendrocytes could be distinguished by cell morphology: OPCs only possessed processes and myelinating oligodendrocytes possessed sheaths. Animal ages and corresponding OPC activity: 2 dpf (when OPCs are born, and migrate), 3 dpf (when many OPCs initiate differentiation), 5 dpf (when many OPCs complete differentiation.)

#### Alignment rate and colocalization ratio

Images of fixed tissue were analyzed in Imaris. “Surface” function was used to create a cell surface (Smooth and Surface Grain Size = 0.0850 μm; Diameter of Largest Sphere = 5 μm; Threshold between 900-1200; Number of Voxels above 100). “Spot” function was used to generate Spot_postsynaptic puncta_ (Estimated XY Diameter = 0.35 μm; Estimated Z Diameter = 0.5 μm; Background subtraction; quality above 2500 to 2800). Then the cell surface was used as a mask to generate Spot_postsynaptic puncta in OPCs_, which was further verified by crosschecking its overlap with Spot_postsynaptic puncta_. Spot_postsynaptic puncta in OPCs_ was analyzed for puncta number or density. To examine the alignment of MAGUK (Gephyrin) and Synapsin, we used “Spot” function to generate Spot_Synapsin puncta_. By determining the proximity Spot_postsynaptic puncta in OPCs_ to Spot_Synapsin puncta_ using MATLAB-enabled “Colocalize Spots” function (<0.1 μm), we could identify postsynaptic puncta in OPCs that aligned with Synapsin and calculate the alignment rate.

For colocalization of PSD-95-GFP (or GFP-Gephyrin) and MAGUK (or Gephyrin) antibody staining, “Surface” function was used to create a GFP surface(Smooth and Surface Grain Size = 0.0850 μm; Diameter of Largest Sphere = 6.5 μm; Threshold between 10000-20000; Number of Voxels above 100). We used the Imaris “Coloc” function to generate a coloc channel based on the overlapping pixels between GFP and MAGUK (Gephyrin) channel and generated a coloc surface (Smooth and Surface Grain Size = 0.0850 μm; Diameter of Largest Sphere = 6.5 μm; Threshold between 3000-12000; Number of Voxels above 100). We calculated the % of GFP that colocalize with antibody staining by dividing the coloc surface volume by the GFP surface volume.

#### Postsynaptic puncta identification and analysis from *in vivo* imaging

Live Images of PSD-95-GFP and GFP-Gephyrin were processed in ImageJ, and synaptic puncta were identified and tracked with Trackmate^73^. In Trackmate, parameters were set as follows: LOG mode, diameter 0.75 μm, Quality>100, contrast>0.19 and signal noise >0.35, LAP linking with 0.5 μm distance and 2 max gap frames allowed were applied. The identified puncta at each frame were then manually examined to remove poor quality tracks. Neighboring puncta may sometimes be incorrectly linked between timelapses; therefore, manual examinations were done to fix these link errors. The data output was analyzed in MATLAB with custom scripts to calculate puncta number per cell at each timepoint, puncta duration and diameter. To generate figures of mean and standard deviation as shade, we used a modified script (https://www.mathworks.com/matlabcentral/fileexchange/29534-stdshade). To analyze the puncta number during OPC differentiation, it was designated as time 0 when myelinating oligodendrocyte’s last process disappeared and all process transformed into sheaths. In MATLAB, this information is used to divide the puncta number into before and after time 0 for each cell. The traces were generated and realigned based on time 0. To analyze the enrichment of puncta in differentiating oligodendrocytes, the puncta number in sheaths and processes were divided by the corresponding areas and normalized.

#### Hotspot analysis

Heatmaps of puncta over time were generated using the output data with Heatscatter (https://www.mathworks.com/matlabcentral/fileexchange/47165-heatscatter-plot-for-variables-x-and-y). With a sampling volume of 1 x 1 x 4 μm, regions with frequency higher than three standard deviations were identified as hotspots using custom MATLAB scripts, which also measured the distance between the center of hotspots and future sheaths. The same set of puncta were used to generate the convex hull for GFP-Gephyrin. In time-lapse imaging that encompassed both OPC and oligodendrocyte stages, we used the first nascent sheath appearance (time 0) to separate OPC stage and oligodendrocyte stage. It takes up to 5 hours for an oligodendrocyte to complete differentiation^36^; therefore, we identified where sheaths formed approximately 6 hours after time 0. We grouped the GFP-Gephyrin puncta into the ones in hotspot regions (PIH) and outside hotspot regions (POH). To choose 30 representative points outside hotspots in an OPC, we used Trackmate to convert OPC processes to points and clustered the points outside hotspot regions into 30 groups (nonhotspots) using Lloyd’s algorithm in MATLAB. The same set of points were used to generate the convex hull for OPC processes.

The distances between an object (hotspots, nonhotspots, PIH, and POH) and myelin sheaths were measured using custom MATLAB scripts. To examine how such a spot predicted myelin sheaths, we analyzed the frequency distribution of spot-sheath distances, and calculated the percentage of distances that were less than 1 μm. To control for the myelin sheath prediction by GFP-Gephyrin puncta, We calculated the percentage of these puncta that were within 1 μm from myelin sheaths. In comparison, we calculated the random chance by dividing the volume occupied by myelin sheaths by the entire volume occupied by OPCs prior to differentiation. To estimate the myelin sheath volume, the sheath length and width were measured and the volume was calculated as pi*length*(width/2)^2^. To estimate the OPC volume over time, we calculated the volume of the shape generated from process-converted points using Delaunay triangulation.

#### Ca^2+^ events analysis

myrGCaMP6s images of z-stacks were maximum projected and stabilized using “Image Stabilizer.” The images were assigned with random names using “blindr” (https://github.com/U8NWXD/blindr) before analysis and reassigned with the original names after analysis. The regions of Ca^2+^ events (ROI) were identified by “GECIquant”^74^ first and then manually examined frame by frame. The intensity and area of each ROI were exported and processed in MATLAB through custom scripts. Briefly, ΔF/F_average_ (ΔF=F-F_average_) was calculated to generate traces and to identify intensity peaks using “findpeaks” in MATLAB with filtering parameters (3 standard deviations above average, prominence larger than 40% of the baseline, and peak area larger than 2.5). Rare elevated Ca^2+^ signal for extended periods of time (longer than 50 seconds) were excluded. Peaks were manually verified against the image. Several Ca^2+^ characteristics were measured: peak amplitude, peak duration, Ca^2+^ MD number, Ca^2+^ MD area, and frequency of Ca^2+^ activity. A region with multiple OPCs was analyzed for TeNTlc and TeNT experiment, while single OPCs were analyzed for the rest of experiments. To examine the proximity of synaptophysin-RFP to Ca^2+^ MD, Ca^2+^ MDs were identified as above, and their locations were first compared to synaptophysin-RFP manually plane by plane and then verified by Imaris. In Imaris, using “Spot” and “Surface” functions, we generated synaptophysin-RFP puncta and OPC surface, with which we identified the puncta that were close to OPC surfaces using a MATLAB-enabled “Spot-to-Surface” function (0.5 μm threshold). These puncta were manually checked against Ca^2+^ MDs identified through morphological landmarks in Imaris. The distance between Ca^2+^ and future sheaths were measured in MATLAB. In TeNTlc experiments, the number of axons expressing TeNTlc-tdtomato was counted in each image. If the number was less than 4, the corresponding image was grouped as “neighboring few TeNTlc axons”; otherwise, the image was grouped as “neighboring many TeNTlc axons”. The distances between Ca^2+^ MDs and future myelin sheaths were measured using custom MATLAB scripts. To choose 30 representative points outside MDs in an OPC, we used Trackmate to convert OPC processes to points and clustered the points outside MD regions into 30 groups (nonMD) using Lloyd’s algorithm in MATLAB. To examine how MDs or nonMDs predicted myelin sheaths, we analyzed the frequency distribution of MD-sheath and nonMD-sheath distances, and calculated the percentage of distances that were less than 1 μm. To examine the Ca^2+^ activity in cell-specific knockdown targeted OPCs, we refined MDs to the mScarlet-labeled processes in order to analyze only the targeted cells.

#### Oligodendrocyte and myelin sheath analysis

To analyze myelin sheath length and number, we used ImageJ ROI manager to trace all sheaths of sparsely labeled oligodendrocytes between segments 1 and 15 in spinal cord. The number of oligodendrocytes analyzed per fish was limited to 3 to ensure even weight among individual fish. Sheath traces and the corresponding images were blinded using “File Name Encrypter” in ImageJ. The sheath length and number were then analyzed in MATLAB with custom scripts and matched with the original genotype following analysis. To calculate the myelinating oligodendrocyte %, in each imaging experiment, the number of oligodendrocytes from one genotype was counted and divided to the total number of cells (both oligodendrocytes and OPCs). In measuring hotspot-sheath distances, we identified the stable myelin sheaths approximately 6 hours after time 0 when processes without terminal sheaths disappeared and all sheaths of oligodendrocytes formed. The transient sheaths were identified similarly to a previous study^47^: larger than 1.5 μm in length and 200% wider than the connected processes. In order to measure the nearest distance between dorsal myelinated axons with EM and IHC images, the center of these axons were identified and a nearest point search was performed using a customized MATLAB script.

#### Statistics

Most data are shown as mean1’±1’SEM., with individual data points shown as circles. We conducted Shapiro-Wilk’s test and Levene’s test to verify data normality and homoscedasticity, respectively, before performing two-sided *t*-test for unpaired groups, paired or ratio-paired *t*-test for paired groups, and one-way ANOVA test for more than 3 groups. If the data did not pass normality tests, we used Mann-Whitney test for unpaired groups, Wilcoxon matched-pairs signed rank test for paired groups, Friedman test for repeated measures of more than 3 groups,

and Kruskal-Wallis test for more than 3 groups. These analyses were performed either in GraphPad Prism or MATLAB. The significance and statistical analyses employed were indicated in figure legends. Animal and cell number used for each experiment, t-values and df for*t*-tests, and *F* values and df for ANOVA were indicated in figure legends. Data collection was not performed blindly due to the experimental conditions, but analyses were performed blindly as indicated. Sample size was chosen based on standard deviation and effect (http://www.biomath.info/power/ttest.htm). Sex is not yet determined in zebrafish at the ages employed in our study and so cannot be considered as a biological variable.

#### Reporting summary

Further information on research design is available in the Nature Research Reporting Summary linked to this manuscript.

#### Data availability

All research materials are available upon request. Source data are provided with this paper.

#### Code availability

All custom scripts are available upon request.

## Notes

### Competing Interest Statement

The authors have declared no competing interest.

### Summary of Updates

New evidences to substantiate our model

