## Extended Figures for "Synapses and Ca^2+^ activity in oligodendrocyte precursor cells can predict where myelin sheaths form"

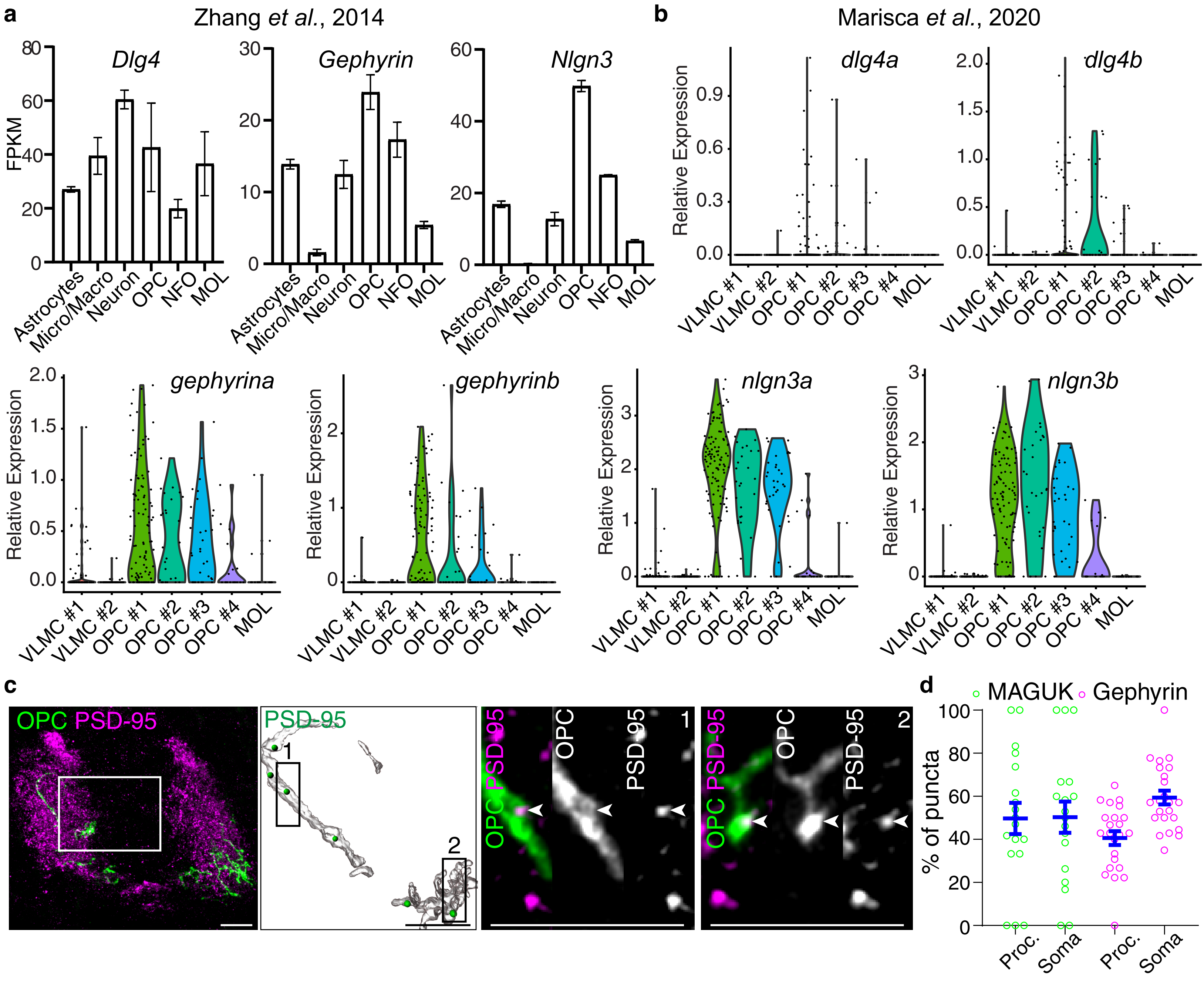


### Extended Data Figure 1

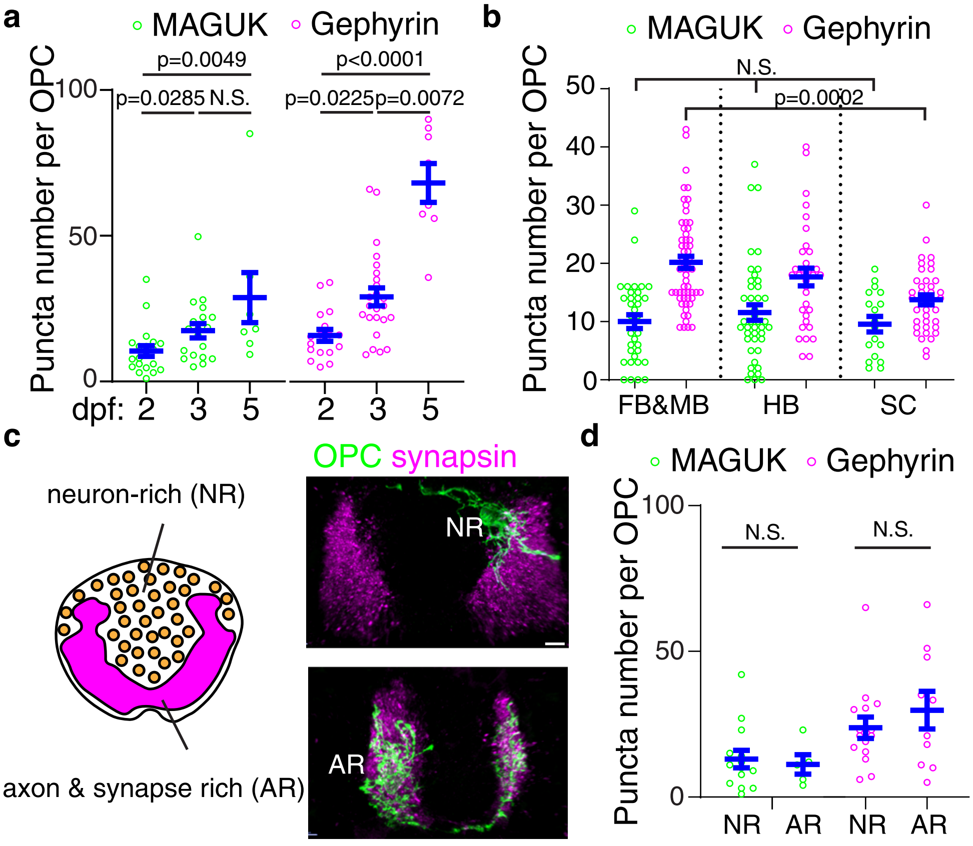


### Extended Data Figure 2

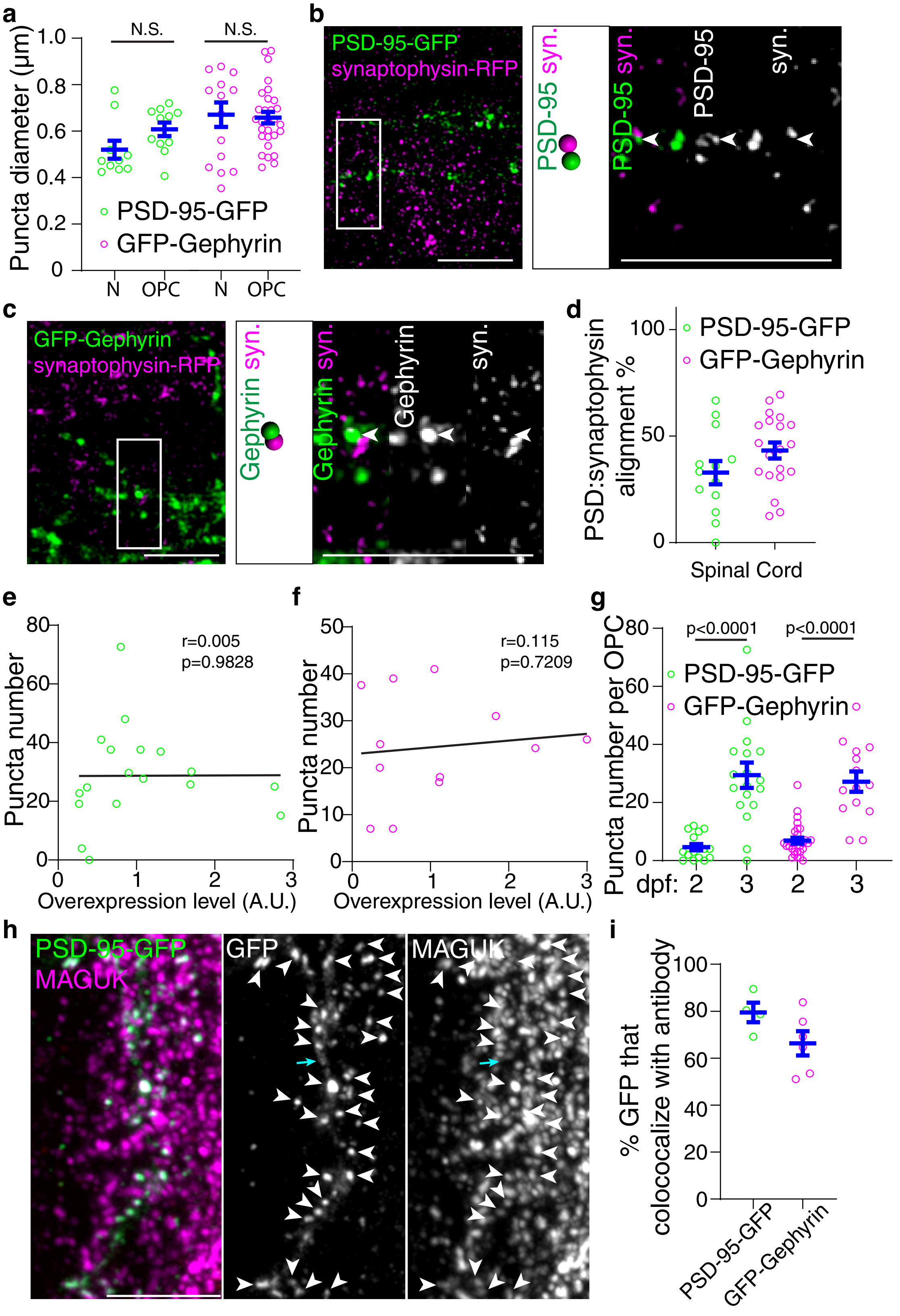


### Extended Data Figure 3

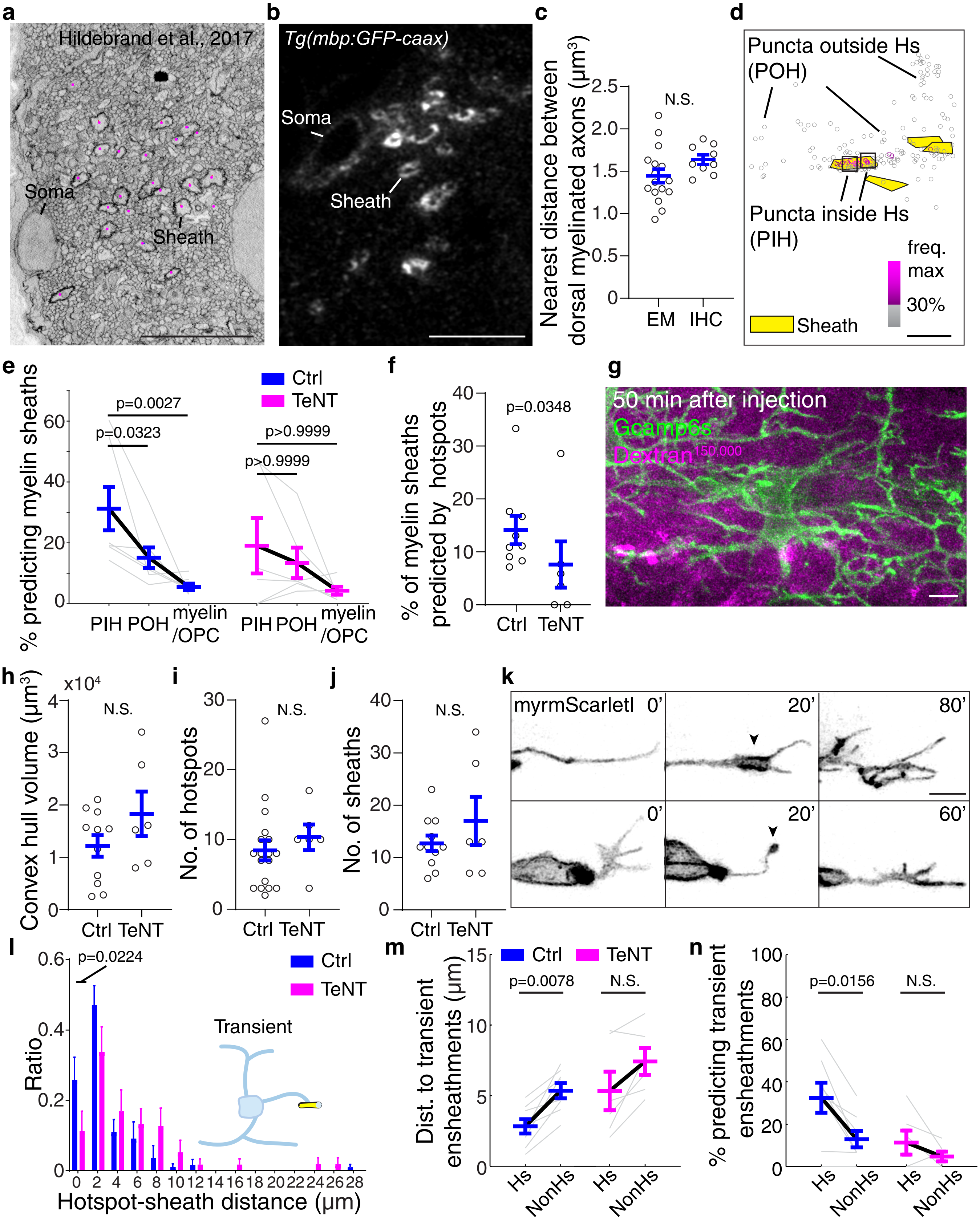


Extended Data Figure 4

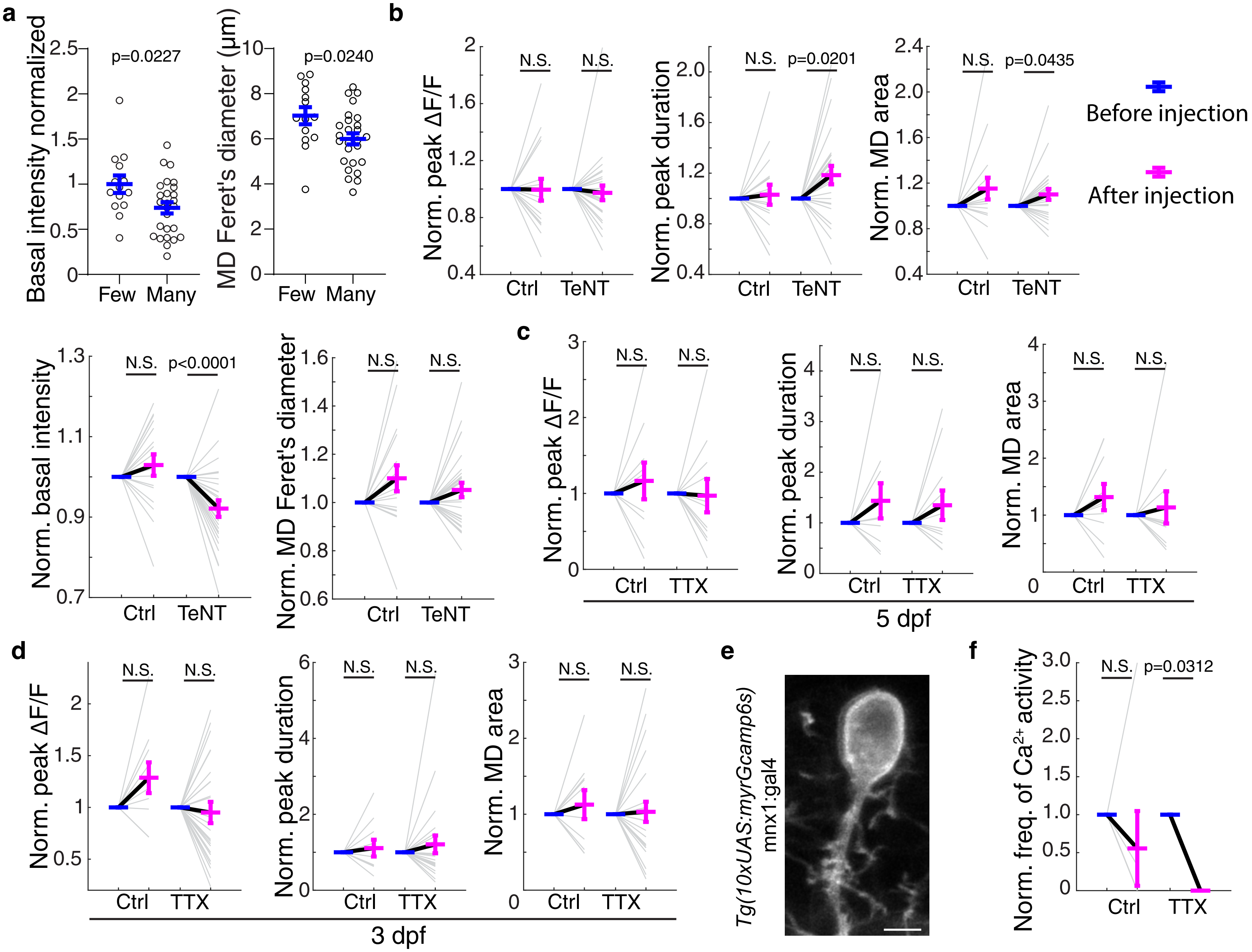


### Extended Data Figure 5

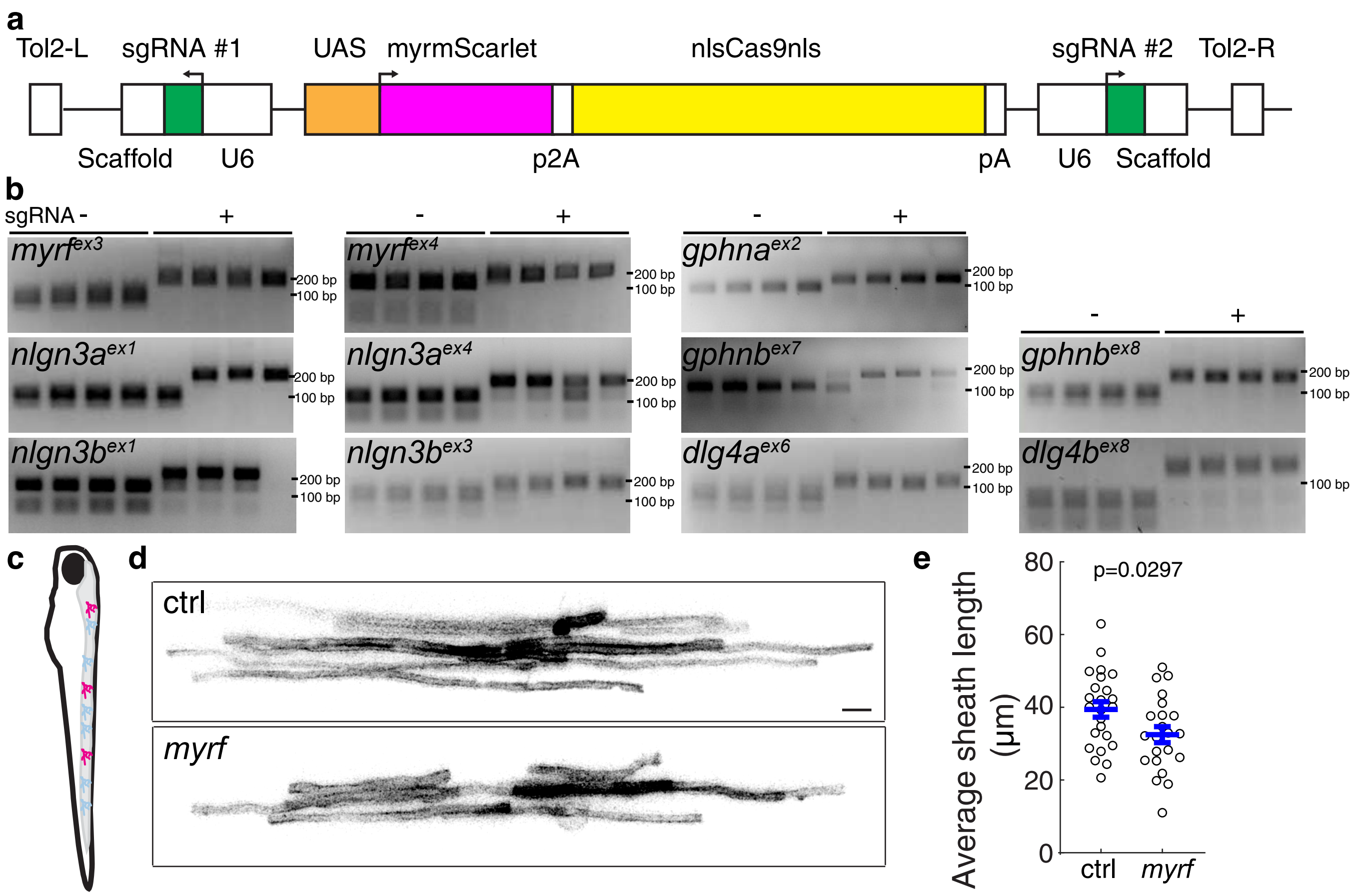


### Extended Data Figure 6

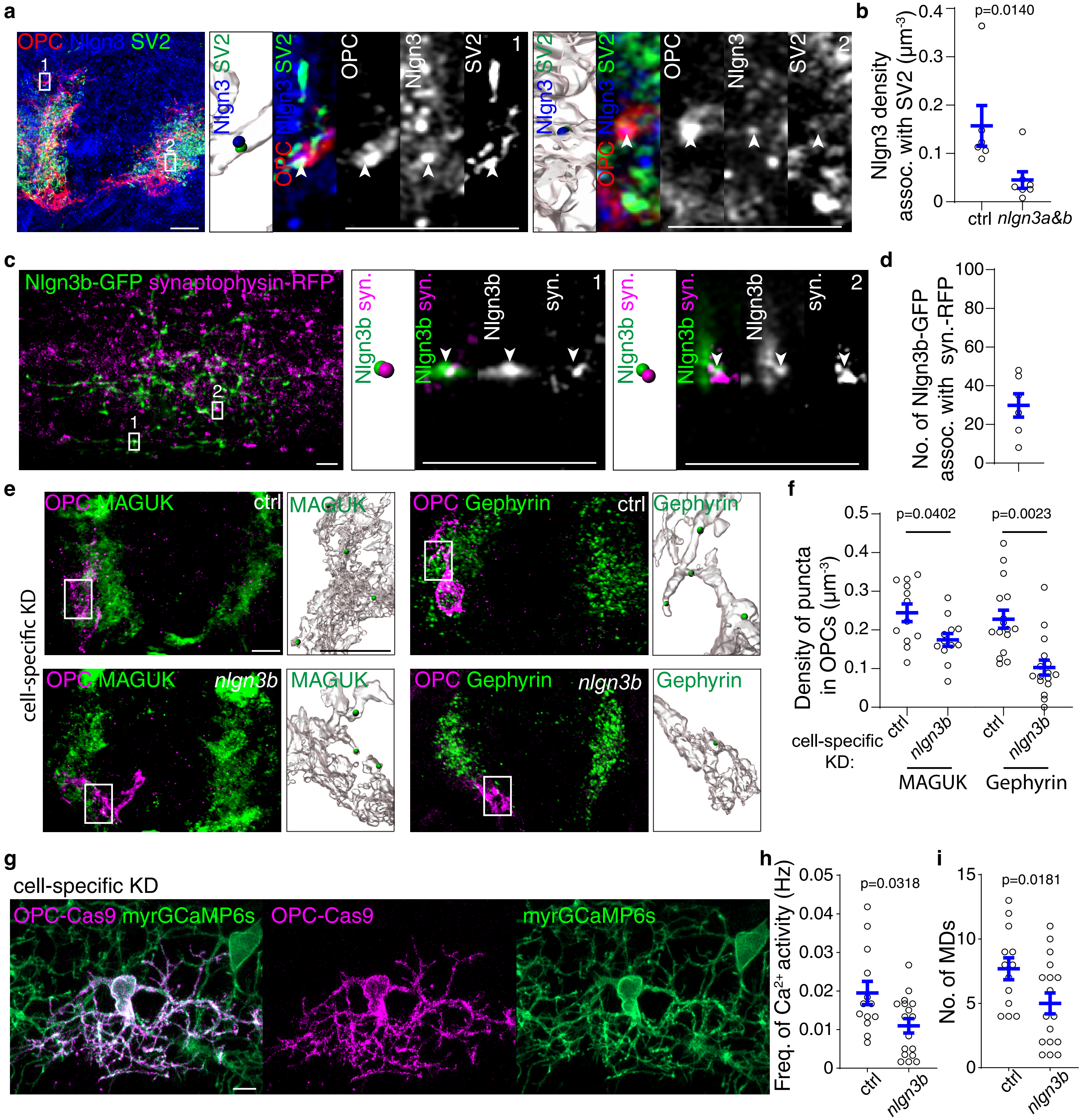


### Extended Data Figure 7

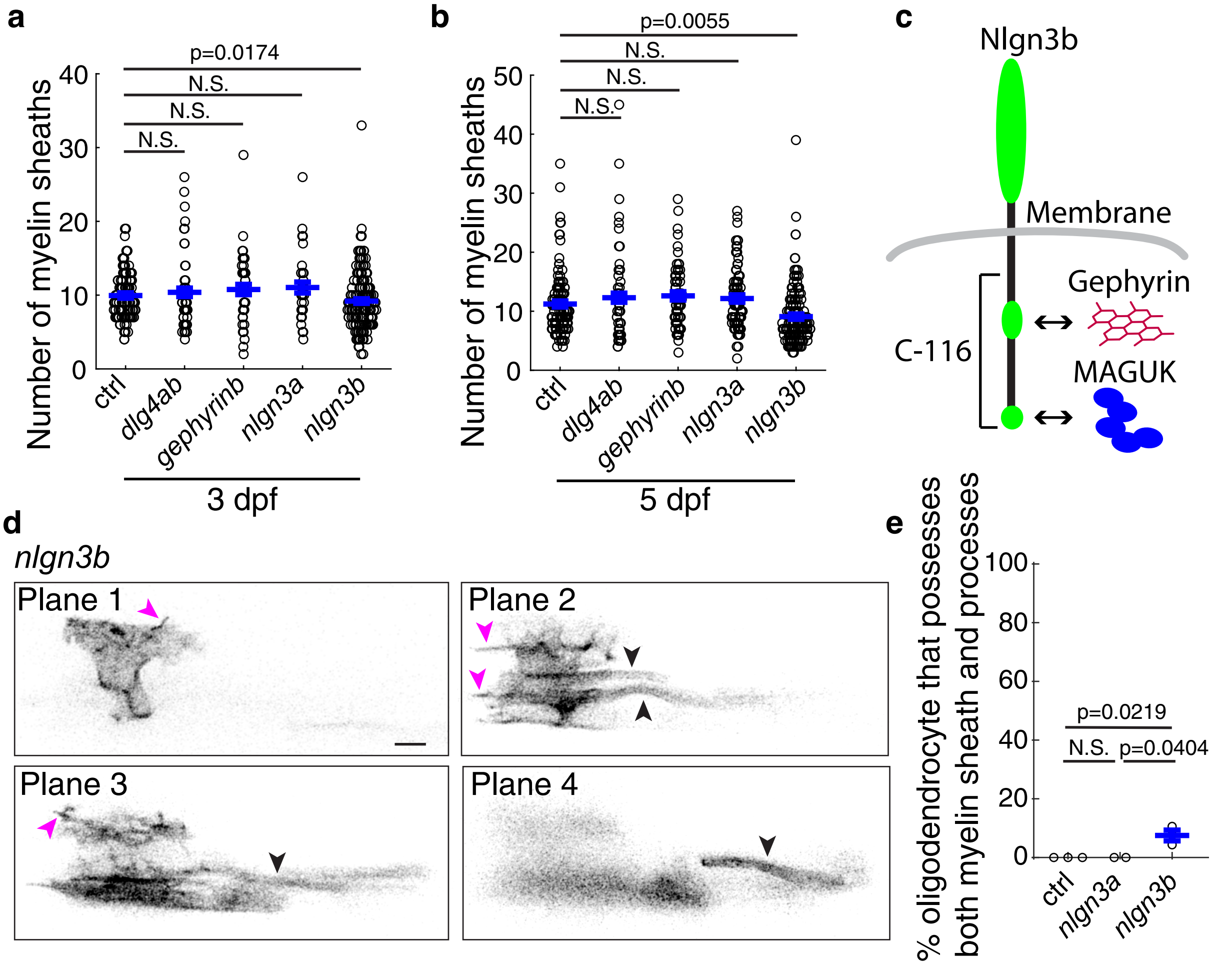


### Extended Data Figure 8
